# Extracellular RNA moves from the glomerulus to the renal tubule

**DOI:** 10.1101/2021.06.15.448584

**Authors:** Robert W Hunter, Sujai Kumar, Richard JM Coward, Amy H Buck, James W Dear

## Abstract

There is a wealth of indirect evidence that extracellular RNA (exRNA) signalling can regulate renal tubular epithelial cell function. However, the physiological importance of this signalling is uncertain. We sought to determine the extent of extracellular RNA transfer between cells in a healthy kidney. We tested the hypothesis that RNA travels from glomerular podocytes to renal tubular epithelial cells.

We developed a method to track exRNA in the kidney using SLAMseq (SH-linked alkylation for the metabolic sequencing of RNA in tissue). We crossed podocin-Cre mice with floxed-stop-UPRT mice to express recombinant uracil phosphoribosyl transferase (UPRT) in podocytes. Mice were injected with the modified nucleobase 4-thiouracil, which is incorporated into nascent RNA with high efficiency only in UPRT-expressing cells. We harvested glomeruli or tubular cells, extracted RNA and prepared libraries for SLAMseq, in which sites of mRNA labelling with 4-thiouracil are detected as T>C conversions in 3’UTRs.

In glomeruli, we detected labelling of known podocyte genes but not of genes known to be restricted to endothelial, renal tubular or white blood cells. Setting a false-discovery rate of 1%, the proportion of genes deemed to be labelled with high confidence was 7.1% (95% confidence interval 6.8 – 7.4%) in 4TU-treated podocyte-UPRT mice, 2.5% (2.3 – 2.7%) in Cre-negative controls and 1.0% (0.9 – 1.1%) in 4TU-naïve controls.

In tubular cells, we detected a small but statistically significant increase in RNA labelling in podocyte-UPRT mice compared to Cre-negative controls (p = 7.4 × 10^−16^ in a zero-inflated Poisson regression model). We conclude that RNA is transferred from podocytes to renal tubular epithelial cells *in vivo* under physiological conditions. Our model provides the opportunity to explore the consequences of this novel signalling pathway in health and kidney disease.

## Introduction

A wealth of indirect evidence suggests that extracellular RNA (exRNA) signalling can regulate renal tubular epithelial cell function.^1–3^ However, this evidence is largely derived from experiments which may not represent *in vivo* physiology because they were conducted *in vitro* or in injury models, or because they relied on single candidate microRNAs being delivered in or knocked out of extracellular vesicles.^e.g. 4–6^ Therefore, we do not know the extent of any exRNA transfer between kidney cells within a healthy kidney.

Indeed, whether exRNA signalling serves a physiological purpose in mammals has been controversial. On the one hand, there are several instances in which exRNA has been shown to regulate mammalian cellular biology^7–13^ and exRNA transport bears hallmarks of a signalling system: exRNA export is selective^14^; uptake by recipient cells is selective^15–17^ and subject to physiological regulation.^18,19^ On the other hand, we do not fully understand mechanisms of RNA uptake or signal transduction and exRNA delivery seems too low to plausibly exert a significant effect on gene expression in some experimental systems.^20^ Historically, experimental approaches have been unable to differentiate between effects mediated directly RNA transferred into recipient cells and effects elicited indirectly by the exposure of recipient cells to exRNA that is not internalised. In the former scenario, exRNA might directly regulate gene expression through canonical sequence-dependent pathways or by activating toll-like receptors;^21^ in the later, exRNA might indirectly perturb gene expression by eliciting a sequence-independent response.

We sought to directly study RNA mobility *in vivo*. We chose to attempt this in the kidney for two reasons. First, the kidney is a tractable system for studying exRNA signalling because the microanatomy and urine flow determine the direction of any exRNA transfer (*from* glomerulus *to* renal tubule). Second, we wanted to determine the potential for exRNA signalling to send injury signals from glomerulus to renal tubule in kidney disease. Renal tubular epithelial cells (rTECs) adopt a pathological phenotype in response to podocyte injury. This is observed in acute podocyte injury (the acute tubular injury that is frequently encountered in adult minimal change disease) and in chronic glomerular disease, which is invariably accompanied by pro-fibrotic, pro-apoptotic, pro-senescent changes in rTECs.^22–24^ This tubular response is observed even in genetic models that deliver a targeted podocyte injury.^25,26^ It is widely assumed that the proteinuria that follows podocyte injury causes rTEC pathology^27^ but it has never been shown that proteinuria is necessary or sufficient to induce rTEC injury *in vivo*. In other contexts exRNA / extracellular vesicle signalling can regulate physiologically important phenomena in rTECs^6,16,18,19^ and the urinary content of podocyte-derived exRNA is elevated 10-100-fold in a range of glomerular diseases including diabetic kidney disease^28^. Therefore, it is at least plausible that exRNA signalling transmits pathological signals from podocyte to rTEC in kidney disease.

We hypothesised that exRNA moves between glomerular podocytes and renal tubular epithelial cells in the mammalian kidney. We tested this directly in mice by specifically labelling podocyte RNA, and then looking for the labelled RNA in rTECs. We used a sequence-based metabolic labelling approach, SLAMseq^29^, to realise podocyte-specific RNA labelling with 4-thiouracil. We provide evidence that RNA is transferred from podocytes to renal tubular cells *in vivo*.

## Methods

### Transgenic mice & animal husbandry

All experimental procedures were performed under UK Home Office licence in accordance with the Animals (Scientific Procedures) Act, 1986 and after review and approval by local veterinary surgeons. Mice had free access to water and a standard RM1 diet (Special Diet Services, Witham, UK) and were maintained on a 12 h–12 h light–dark cycle.

Floxed-stop-UPRT mice, B6;D2-Tg(CAG-GFP,-Uprt)985Cdoe/J^30^, and mTmG mice^31^, Gt(ROSA)26Sor^tm4(ACTB-tdTomato,-EGFP)Luo^/J, were purchased from Jackson laboratories. Podocin-Cre mice, Tg(Nphs2-cre)1Nagy^32^, were re-derived in Edinburgh using sperm from the Bristol line (RJMC’s laboratory). C57BL/6JCrl wild-type mice were purchased from Charles River laboratories.

Floxed-stop-UPRT mice were maintained through homozygous-homozygous crosses. Podocin-Cre mice were maintained through crosses between hemizygous males and wild-type females (to avoid any deleterious effects of germline Cre expression in the female). Podocin-UPRT mice were generated through crosses between podocin-Cre^Tg/-^ males and floxed-stop-UPRT^Tg/Tg^ females. Mice of both sexes were used in experiments.

### Genotyping

Genomic DNA was extracted from ear punch biopsies using the “HotSHOT”method.^33^ Punch biopsies were digested in 75 microL alkaline lysis reagent (25 mM NaOH, 0.2 mM EDTA) at 95 °C for 30 minutes and then cooled on ice. 75 microL of neutralisation buffer (40 mM Tris-HCl) was added and samples centrifuged briefly to pellet large debris. 1 microL of the supernatant was used as a PCR template.

PCR was performed using Taq polymerase (Invitrogen; 0.04 units / ml) with 0.2 mM dNTPs and 1.5 mM Mg^2+^. Forward primers were used at 0.5 fM; reverse primers at 0.5 fM for double-primer reactions or 0.25 fM each for triple-primer reactions. Primer sequences are given in Table 1. Reactions were carried out using a “touchdown” cycling program: the annealing temperature was reduced by 1 °C per cycle – from 72 °C to 60 °C – for an initial 12 cycles and held constant at 58 °C for a final 24 cycles.

**Table 1.**
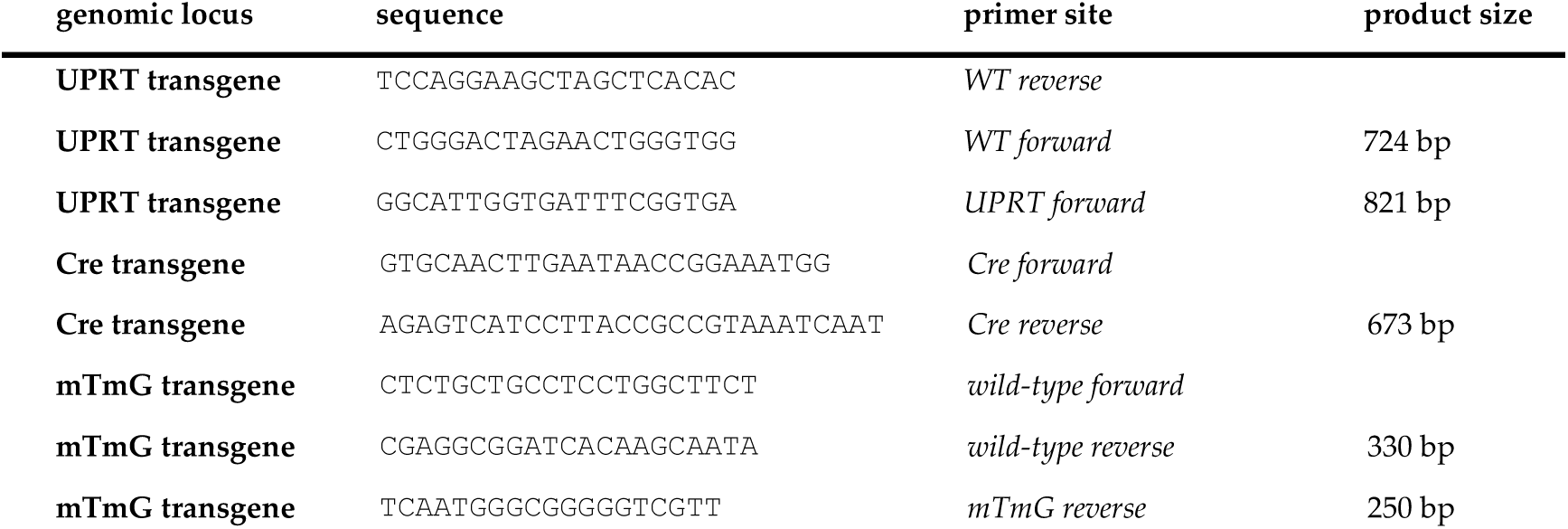
Genotyping primers.

### Immunofluorescence

Indirect immunofluorescent detection of target antigens was performed using a Leica BOND-MAX TM robot after an antigen retrieval step (citrate buffer, pH6 for 20 minutes). The binding sites of HRP-conjugated secondary antibodies were detected with Red Opal (1:100, AKOYA Biosciences) and a DAPI counterstain was applied (1:1000). Antibodies are listed in Table 2.

**Table 2.** Antibodies used for immunoblot and immunofluorescence.

| antibody | source | application | dilution |
| --- | --- | --- | --- |
| nephrin | R&D systems (AF3159) | immunoblot (primary) | 1:5,000 |
| HA tag | CST (C29F4) | immunoblot (primary) | 1:1,200 |
| beta actin | CST (13E5) | immunoblot (primary) | 1:1,200 |
| protein-G HRP | Abcam (ab7460) | immunoblot (secondar) | 1:1,000 |
| anti-goat-HRP | Santa Cruz (sc-2354) | immunoblot(secondary) | 1:1,000 |
| podocin | Abcam (ab50339) | immunofluorescence (primary) | 1:10,000 |
| HA tag | CST (C29F4) | immunofluorescence (primary) | 1:10,000 |
| anti-rabbit-HRP | Abcam (ab7171) | immunofluorescence (secondary) | 1:500 |

### Fluorescence microscopy

Specimens from the mTmG mouse were prepared for fluorescence microscopy by cryopreservation.^31^ After CO_2_ euthanasia, mice were perfused with cold, filtered 0.9% NaCl through the left ventricle. Kidneys were decapsulated, bisected and immersed in 4% methanol-free PFA (Thermo 28908) in PBS for 2 hours in the dark at 4 °C. They were twice washed with PBS before being immersed in 18% sucrose overnight in the dark at 4 °C. The samples were flash-frozen in OCT and stored at –80 °C until sectioned. 7 μm sections were washed thrice with PBS and stained with DAPI. Epifluorescence microscopy was performed on an Axioscan SlideScanner; confocal microscopy on a Zeiss LSM 710.

### Immunoblot

Whole kidneys were homogenised in 250 mM sucrose / 20 mM triethanolamine with protease inhibitors (1% Merck Protease Inhibitor Cocktail III). The homogenate was cleared of large debris by centrifugation at 4000 *g* for 15 mins at 4 °C. Glomerular samples were lysed in RIPA buffer, with protease inhibitors as above.

Samples were prepared for SDS-PAGE by mixing with 2x tris-glycine SDS sample buffer (Invitrogen) and DTT (final concentration 50 mM) and then heating to 70 °C for 15 mins. SDS-PAGE was carried out using Novex WedgeWell 4-12% Tris-glycine gels. Gels were blotted onto PVDF by wet transfer in Tris-glycine buffer with 20% methanol. Membranes were then washed in 0.2% Tween in PBS and blocked in 5% (w/v) milk powder before being incubated with primary antibodies at 4 °C overnight. After three washes in 0.2% Tween in PBS, membranes were incubated with HRP-conjugated secondary antibody for 60 mins at room temperature before being washed thrice again. Antibodies are listed in Table 2. HRP signal was detected using ECL reagent and a LICOR Odyssey imager.

### Isolation of glomeruli

Glomeruli were isolated after perfusion with magnetic nanobeads.^34^ 4.5 micron tosylactivated superparamagnetic Dynabeads (Thermo 14013) were first inactivated in 400 microL batches. They were washed twice with 1% BSA in PBS and then re-suspended in 1 ml 0.2M Tris-Cl / 0.1% BSA, pH 8.0 and incubated overnight on a rocking platform at room temperature. The following morning they were resuspended in HBSS (Thermo 14175-095: Ca^2+^ and Mg^2+^-free) at a concentration of ∼8 × 10^7^ (= 200 microL) beads per 10 ml.

Under terminal isofluorane anaesthesia, mice were perfused through the left ventricle with 20 ml HBSS and then with 8 × 10^7^ Dynabeads in 10 ml HBSS. Kidneys were dissected into ice-cold HBSS and then diced, on ice, into ∼1 mm^3^ chunks. They were digested in 1.5 ml of 1 mg/ml collagenase A (Roche 10103578001) / 100 U/ml DNAasI (Invitrogen) in HBSS (14025-092: containing Ca^2+^ and Mg^2+^). They were incubated for 30 mins at 37 °C on a rotating rack; this step was repeated if large chunks of kidneys remained. The supernatant was then passed through a 100 micron cell strainer, which should not retain intact glomeruli. The residual kidney chunks were resuspended in digestion buffer and digested for a further 20 minutes at 37 °C; this digest was passed through the same 100 micron cell strainer. The pooled filtrates were passed again through a 100 micron cell strainer (and washed through with HBSS). The glomeruli were gently pelleted by centrifugation at 500 *g* for 5 mins, then re-suspended in 800 microL HBSS and transferred to a 1.5 ml microfuge tube. Glomeruli were concentrated on a magnet and subjected to 2 washes with HBSS.

### Isolation of renal tubular cells

Under terminal isofluorane anaesthesia, mice were perfused through the left ventricle with 20 – 40 ml ice-cold PBS. Kidney were dissected into ice-cold PBS and then diced – on ice – into ∼1 mm^3^ chunks. They were placed into 4 ml digestion buffer (0.425 mg/ml Collagenase V; 0.625 mg/ml Collagenase D; 1 mg/ml Dispase II; 100 units/ml DNaseI in RPMI medium). They were then transferred to MACS tubes (130-096-334) and agitated on the gentleMACS Octo Dissociator before being incubated for 30 mins at 37 °C with gentle agitation. Digestion was quenched with 4 ml ice-cold neutralisation buffer (2 % fetal bovine serum, 1 mM EDTA in Ca^2+^ and Mg^2+^-free PBS). Samples were then passed through 100, then 70 then 40 micron cell strainers, retaining the filtrate each time. Cells were pelleted at 500 *g* for 5 mins and then resuspended in 1 ml RBC lysis buffer (R7757). After 60 seconds, samples were diluted into 15 ml ice-cold neutralisation buffer. Cells were pelleted as before and resuspended in 200 microL neutralisation buffer.

We used Lotus Tetragonolobus Lectin (LTL) to bind to renal proximal tubular epithelial cells.^35,36^ Cells were then incubated with LTL-biotin (VectorLabs B-1325) at 1:100 dilution and incubated at 4 °C for 30 mins. After three washes in neutralisation buffer, cells were incubated with Streptavidin-APC (Thermo 17-4317-82) at 1:400 dilution at 4 °C for 30 mins in the dark. Cells were then washed again thrice.

rTECs were selected as GFP+, LTL+ cells on an a FACSAria II cell sorter. GFP-negative, unstained, unconjugated LTL (VectorLabs L-1320) and streptavidin-negative controls were included in every experimental batch. Cells were collected into PBS and then pelleted at 1000 *g* for 10 mins. Most of the supernatant was aspirated, leaving ∼250 microL; this was then added to 750 microL TRIzol LS and samples were homogenised by vortexing.

### 4-thiouracil treatment

4-thiouracil (Sigma 440736) was dissolved at 200 mg/ml in DMSO and stored in small aliquots at –20°C. On the day of administration, this stock was diluted 1:10 in corn oil to give a 20 mg/ml solution in 90% corn oil / 10% DMSO. Mice received a 20 ml/kg (= 400 mg/kg body weight) intraperitoneal injection at 48, 24 and 12 hours prior to euthanasia.

### RNA extraction and alkylation

RNA was extracted and alkylated as described in the original SLAMseq protocols.^29,37^ Briefly, cell or tissue samples were lysed in a monophasic phenol / guanidine isothiocyanate solution (TRIzol LS for cells; TRIsure for glomeruli or whole kidneys) and then RNA extracted by chloroform extraction and isopropanol precipitation in the presence of 20 mcg glycogen and 0.1 mM DTT. Samples were treated with DNAse (Thermo AM1907) and then cleaned up on silica columns (Zymo R1013), eluting into 15 microL 1 mM DTT. RNA samples were alkylated by treating with 10 mM iodoacetamide in 50 mM sodium phosphate buffer / 50 % DMSO for 50 °C for 15 mins. The reaction was quenched with 20 mM DTT and then RNA samples purified by ethanol precipitation. The integrity and concentration of alkylated RNA samples was assessed by automated capillary electrophoresis (on a LabChip GX Touch). To verify success of the alkylation reaction, 1 mM 4-thiouracil controls were included and diluted 1:10 before being subjected to UV absorbance spectrophotometry (Nanodrop in UV-Vis mode).

### RNA sequencing & SLAMDUNK analysis

Alkylated RNA samples were submitted to Lexogen who prepared libraries for QuantSeq using 50 ng input total RNA and 18 cycles of PCR library amplification. Sequencing was performed on a NextSeq 500 in single read mode using a 75 cycle high output cartridge. QuantSeq is a method of sequencing the 3’ ends of mRNA.^38^ The SLAMDUNK analysis pipeline was used to map and quantify T>C conversions.^29,39^

### Cell-enriched genes in reference dataset

To define a set of known cell-enriched (or “marker” genes) we used published single cell RNAseq data from mouse kidney.^40^ Supplemental table S3 from Park et al. lists the proportion of any given cell type expressing any of a list of genes. We first excluded the “Novel” cell types and lumped together all tubular and white blood cell (WBC) sub-types, setting the % of positive cells as the maximum value for any cell type within each group. We then – arbitrarily – defined a gene as being enriched within a particular cell type if it was present in at least 5% of cells of that type and in no more than 1% of cells of any other type.

### Podocyte injury model (doxorubicin)

Doxorubicin (Adriamycin; Sigma D1515) was dissolved in 0.9% NaCl to give a 3 mg/ml solution and then filter-sterilised. This was administered by tail-vein injection at a dose of 15 micrograms per gram body weight. Control mice received no injection. Urine samples were collected by allowing mice to roam freely in a cage lined with LabSand® at 1000 hrs and again at 1500 hrs on collection days; morning and afternoon urine samples were pooled. Mice were culled by CO_2_ euthanasia. A terminal blood sample was obtained immediately afterwards *via* cardiac puncture.

Serum creatinine was determined using the creatininase/creatinase specific enzymatic method described by Bömer using a commercial kit (Alpha Laboratories Ltd. Eastleigh, UK) adapted for use on a Cobas Fara centrifugal analyser (Roche Diagnostics Ltd, Welwyn Garden City, UK).^41^ Within run-precision was CV <3%.

Serum albumin measurements were determined using a commercial serum albumin kit (Alpha Laboratories Ltd., Eastleigh, UK) adapted for use on Cobas Mira analyser (Roche Diagnostics Ltd, Welwyn Garden City, UK). The measurement of serum albumin is based on its quantitative binding to bromocresol green (BCG). The albumin-BCG-complex absorbs maximally at 578nm, the absorbance being directly proportional to the concentration in the sample. Within-run precision was CV <2.5%.

Urine albumin measurements were determined using a commercial Microalbumin Kit (DiaSys Diagnostics Systems, Germany) adapted for use on a Cobas Mira analyser. The immunoturbidimetric assay was standardised against purified mouse albumin standards (Sigma Chemical Co. Poole, UK) with samples diluted in phosphate buffer saline as appropriate. Within-run precision was CV <5%.

### Data analysis and statistics

Data were analysed in R (version 3.6.1)^42^ using the Tidyverse package (version 1.3.0)^43^. Our code is provided at https://github.com/robertwhunter/podoSLAM.

T>C conversion rate distributions were plotted using the ggridges package (v0.5.3).^44^ Given the distribution of T>C conversion rates, in which the majority of genes had zero T>C conversions, we chose to analyse these data using a zero-inflated Poisson regression model (pscl package, v1.5.5).^45^ (In sensitivity analyses, we also used the Kruskal-Wallis rank sum test and two-way Kolmogorov-Smirnov test to make between-group comparisons; the conclusions were not significantly altered by taking either of these approaches.)

T>C conversion rate data did not follow a normal distribution, limiting the use of the mean as a useful summary statistic. Due to the large proportion of unlabelled reads, the median in many groups was zero and was therefore a meaningless summary statistic. We therefore used quantile 0.8 (i.e. 80^th^ centile) as a summary statistic, as this fell approximately in the middle of the distribution of genes in which the T>C conversion rate was greater than zero. 95% confidence intervals were derived by the bootstrap method, sampling 10,000 times with replacement. We used the R boot package (v.1.3-28) and the “normal” method for constructing confidence intervals; using alternative methods gave near-identical results.^46^

Flow cytometry data were analysed using the flowCore (v1.11.20) and flowWorkspace (v0.5.40) packages.^47,48^ FACS data were normally distributed; between-group comparisons were made by analysis of variance (ANOVA).

## Results

### Determining the genomic locus of the floxed-stop-UPRT transgene

In order to develop a PCR genotyping assay capable of distinguishing between homo- and hemizygosity, we first determined the genomic locus of the floxed-stop-UPRT transgene. Following the method described by Liang et al., we digested genomic DNA with *NcoI* and then treated with T4 DNA ligase in an attempt to circularise the restriction fragments.^49^ From the known transgene architecture, we predicted that this would generate concatemers of ∼1100 bp in addition to ligates of unknown size generated from the 5’ and 3’ ends of the transgene insertion site (Figure 1A).^30,50^ These samples were used as templates in a PCR reaction, with primers facing away from the body of the transgene, so as to amplify a region including flanking genomic sequence (Figure 1A). The PCR products were resolved by gel electrophoresis and then sequenced using the Sanger method. One product contained sequence running into the 3’ end of the transgene from a locus on chromosome 12 (104426100-104425987); a flanking *NcoI* site gave a predicted product size of 712 bp, matching the size of the observed band.

**Figure 1.**
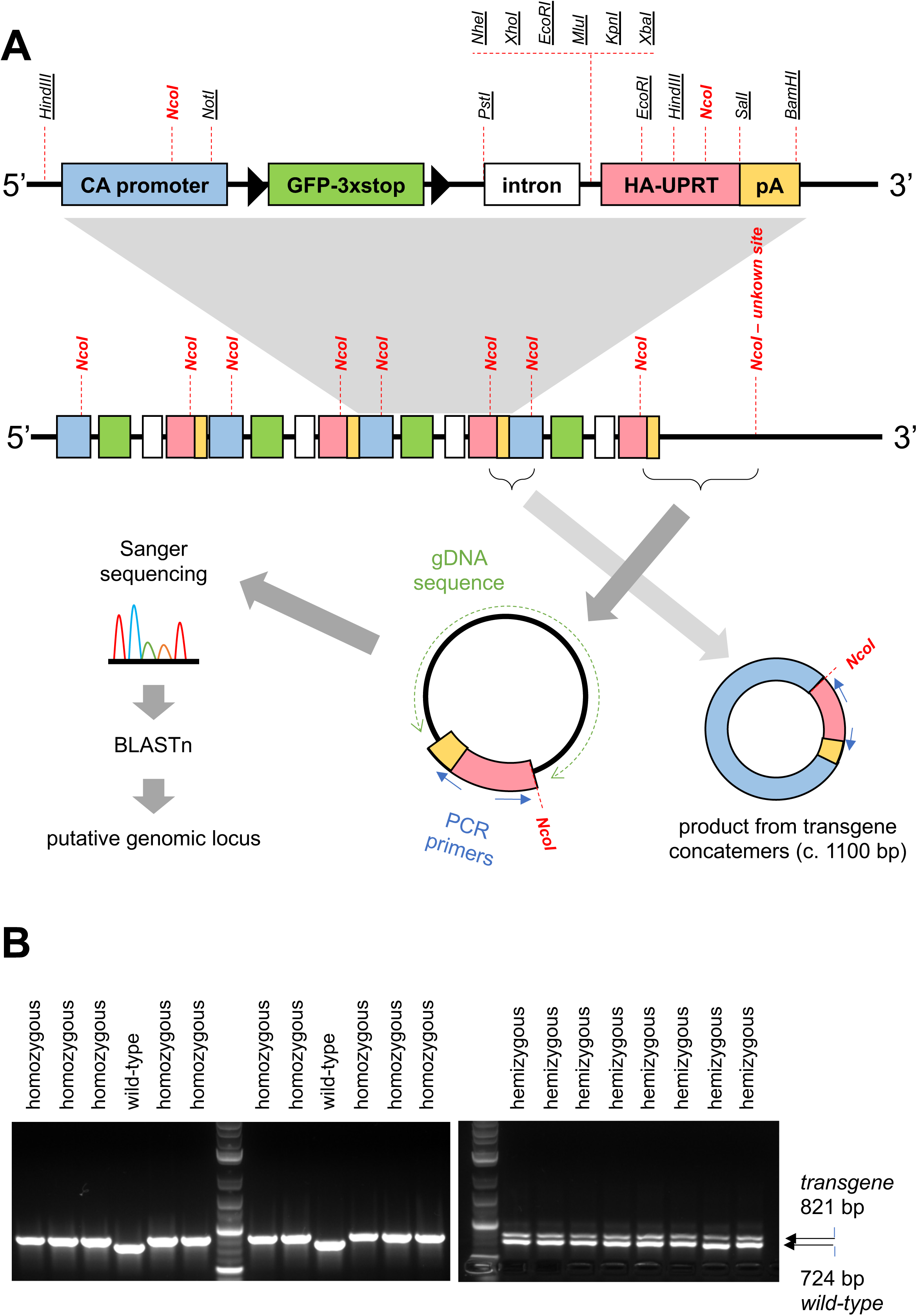
Determining the genomic locus of the floxed-stop-UPRT transgene. **A) Strategy for determining the genomic locus.** The uppermost panel shows the architecture of the floxed-stop-UPRT transgene with approximate location of restriction sites (not to scale). It is likely that this will have integrated as multiple concatemeric copies, as shown below. We digested genomic DNA with *NcoI* and then treated with DNA ligase in an attempt to circularise restriction fragments. These were then used as templates in a PCR reaction, with primers designed to amplify any adjacent stretches of genomic DNA. This strategy revealed a putative genomic locus on chromosome 12. **B) Final genotyping reactions.** We designed a triple-primer genotyping assay with a common reverse primer sited immediately 3’ to the transgene and unique forward primers recognising wild-type or transgene sequence. A 724 bp wild-type band was detected in wild-type and hemizygous mice (but not in homozygous mice); an 821 bp transgene band was detected in hemi- and homozygous mice (but not in wild-types).

We confirmed that this was the single site of transgene insertion, using PCR to amplify an 821 bp product across the 5’ end of the transgene or a 724 bp product across the integration site in the wild-type locus (Table 1; Figure 1B).

### Generation and validation of podocin-UPRT mice

To verify efficient and podocyte-specific Cre recombinase expression, we crossed podocin-Cre with mTmG reporter mice.^31^ In the Cre^+/-^, mTmG^+/-^ offspring, podocytes exhibited green membrane fluorescence whereas all other cell types exhibited red membrane fluorescence, confirming podocyte-restricted Cre expression (Figure 2A; supplemental figure S1).

**Figure 2.**
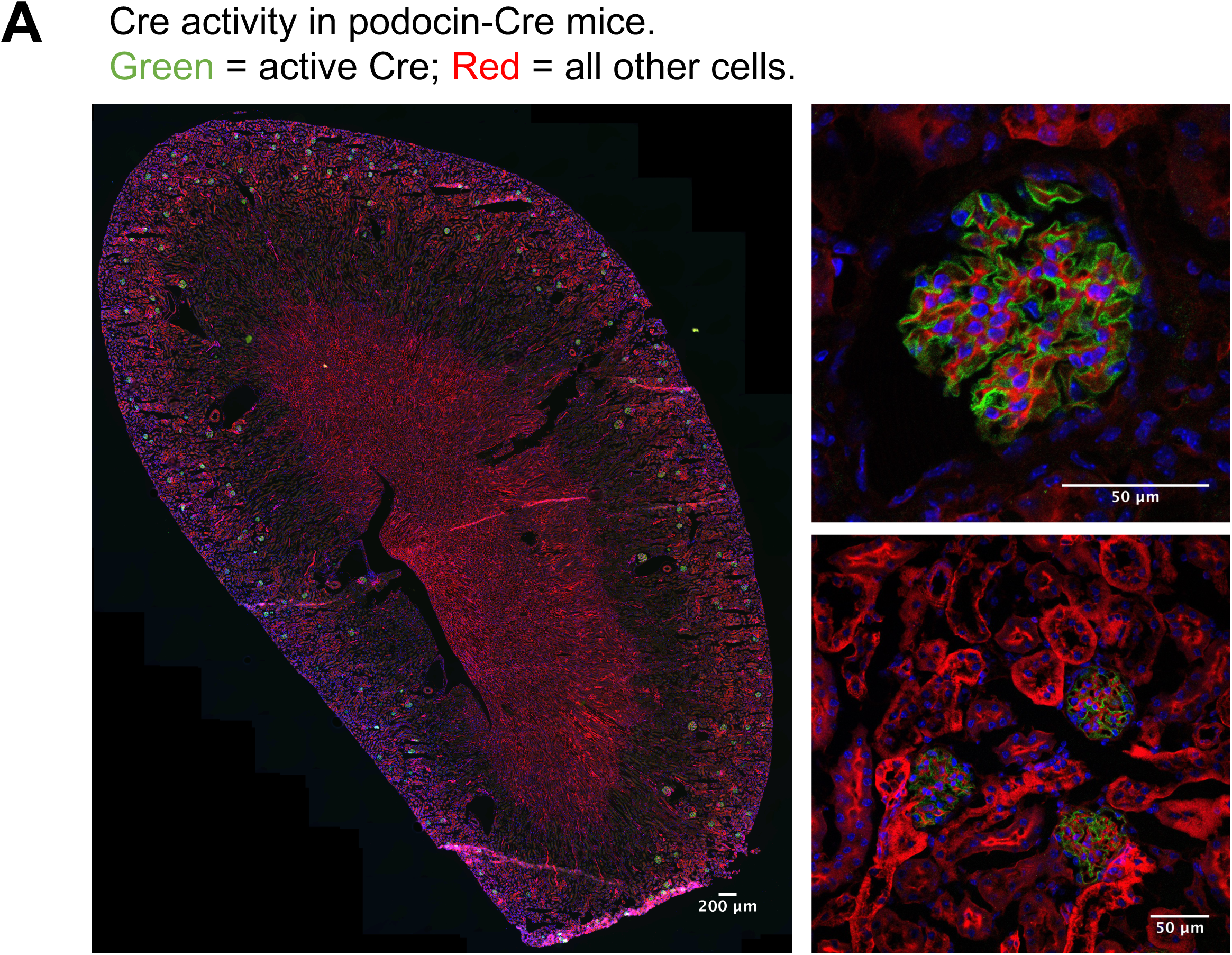
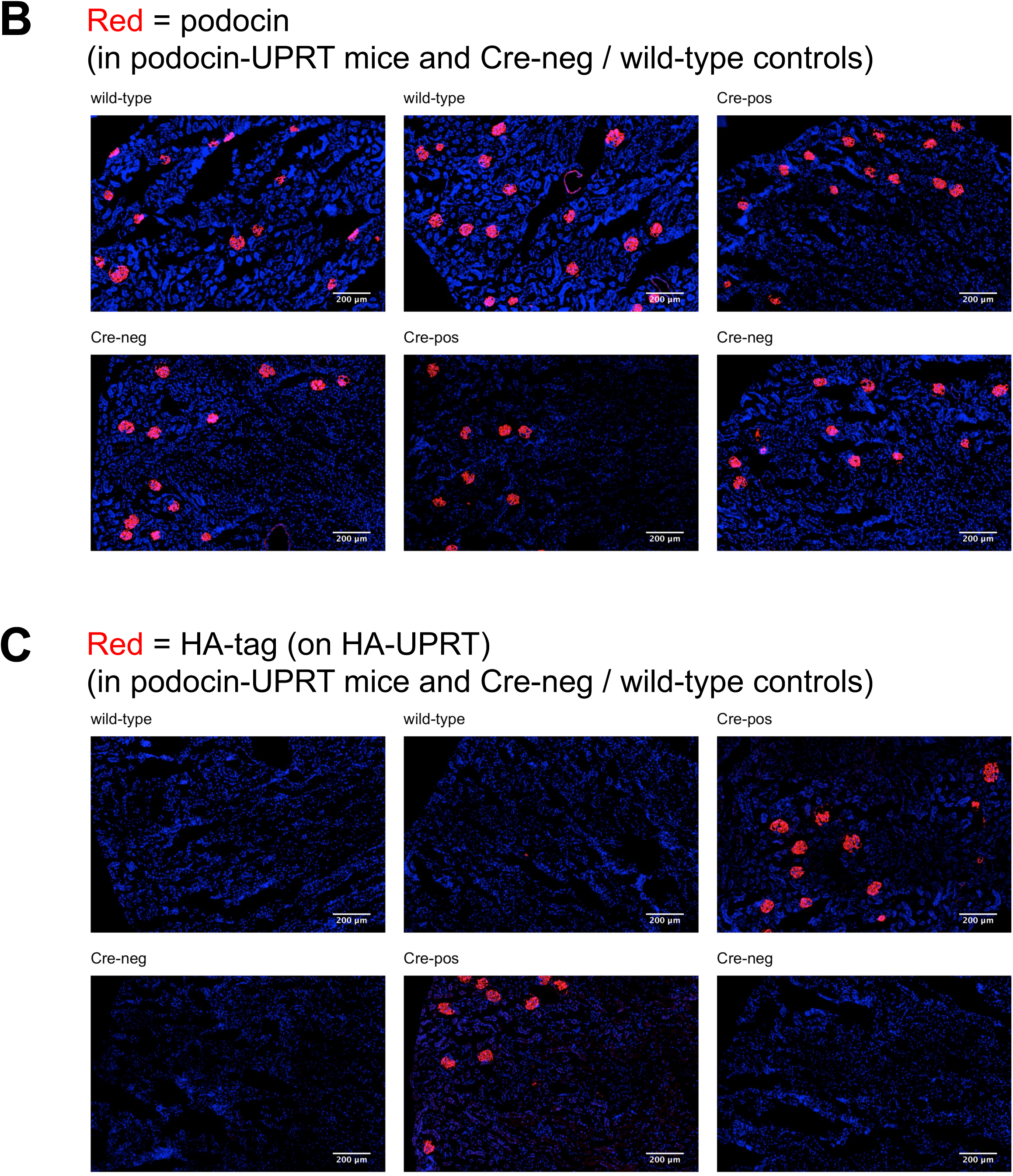
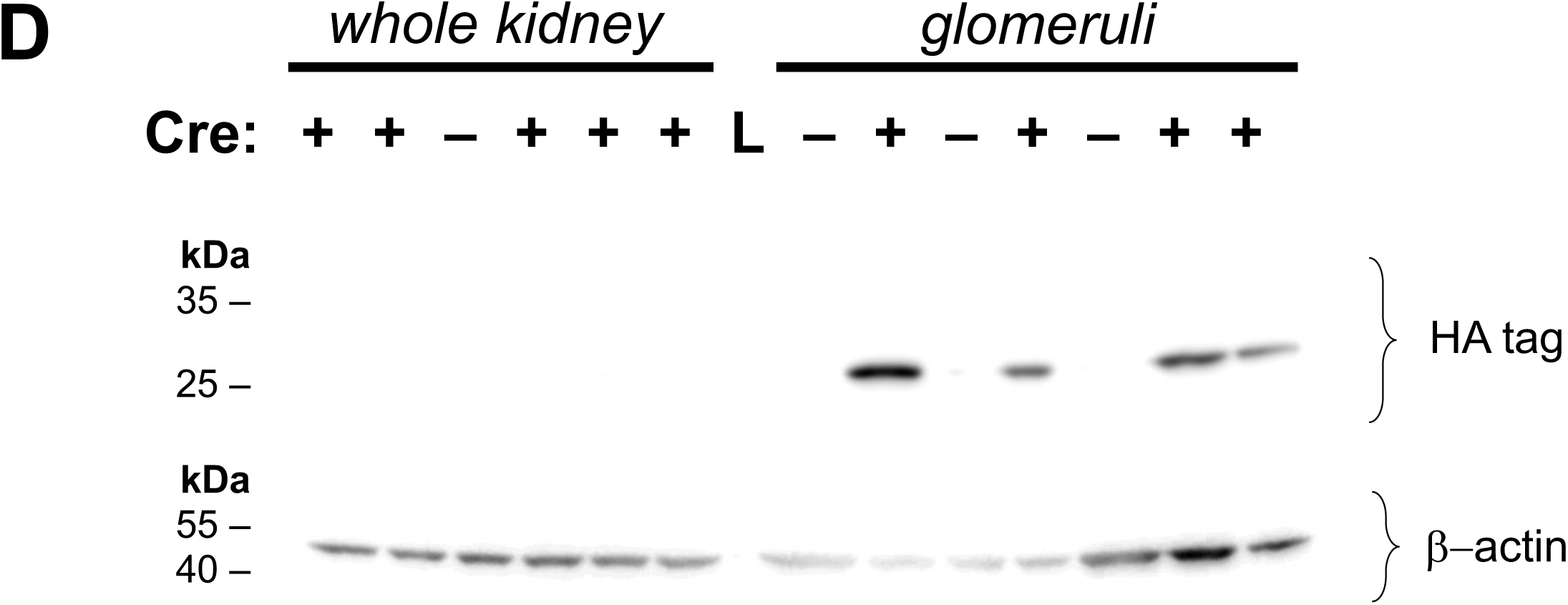
Expression of Cre recombinase and HA-UPRT in podocin-UPRT mice. **A) Site of Cre recombinase expression, reported in mTmG mice.** Podocin-Cre mice were crossed with mTmG mice; kidney sections from the offspring were imaged by fluorescence microscopy. All cells express a red fluorescent membrane protein unless there has been expression of Cre recombinase, in which case expression switches to a green fluorescent protein. GFP expression was observed only in podocytes in Cre^+^ mice. Representative epifluorescent and confocal micrographs are shown. **B) Indirect immunofluorescent detection of podocin. C) Indirect immunofluorescent detection of HA tag.** The haemagglutinin tag (in HA-UPRT) was detected by indirect immunofluorescence in the podocytes of Cre ^Tg/−^ UPRT^Tg/−^ mice but not in Cre^−/−^ or UPRT^−/−^ controls. **D) Detection of HA tag by immunoblot in glomeruli.** Whole kidneys or glomeruli were used to prepare protein for immunoblotting. Kidneys were obtained from podocin-UPRT mice (denoted Cre+) or Cre-negative littermate controls. The lane marked “L” was used to run the ladder. 20 micrograms of protein were loaded per lane. After blotting the membrane was cut horizontally at ∼35 kDa; the upper portion was probed with anti-βactin (expected band at 45 kDa) and the lower portion probed with anti-HA-tag (expected band at 28 kDa). The HA tag was detected by immunoblot in Cre ^Tg/−^ UPRT^Tg/−^ mice but not in Cre^−/−^ or UPRT^−/−^ controls.

Hemizygous podocin-Cre males (Cre^+/-^) were crossed with homozygous floxed-stop-UPRT (UPRT^Tg/Tg^) females to generate podocin-UPRT (Cre^+/-^, UPRT^Tg/-^) and Cre-negative littermate controls. To verify podocyte expression of HA-UPRT, we performed indirect immunofluorescent detection of the HA tag. This tag was present in podocytes in podocyte-UPRT mice (Cre^+/-^, UPRT^Tg/-^) but not in Cre-negative littermate (Cre^-/-^, UPRT^Tg/-^) or wild-type controls (Figure 2B&C). Similarly, immunoblot detection of the HA tag in glomerular preparations gave a strong signal in podocyte-UPRT mice but not in controls (Figure 2D).

### Characterisation of podocin-UPRT mouse

Before embarking on RNA labelling studies, we performed some basic genetic and phenotypic characterisation of podocyte-UPRT mice. Genomic DNA from two podocyte-UPRT mice was analysed by a SNP panel testing 238 alleles (by Transnetyx). ∼90% of tested alleles were from C57BL/6 strain; the majority of the remaining alleles were from the 129 strain (Supplemental table S1).

C57BL/6 strains are often relatively resistant to glomerular injury. Therefore to assess their suitability for modelling podocyte injury / glomerular disease, mice were treated with intravenous doxorubicin (adriamycin).^51^ During the first 8 days after injection, mice exhibited progressive loss of body weight but otherwise exhibited no signs of distress Doxorubicin treatment caused a reduction in serum creatinine and an increase in urinary albumin excretion (Supplemental figure S2).

### Method for labelling podocyte RNA with 4-thiouracil

Mice were labelled with 4-thiouracil (400 mg/kg by IP injection at 48, 24 and 12 hours before cull). Mice were used to provide kidneys either for glomerular samples or for isolation of tubular epithelial cells by FACS (Figure 3; supplemental figure S3). To confirm that our protocol did indeed isolate glomeruli, we tested samples of glomerular preparations and whole kidneys for the presence of known podocyte proteins by immunoblot (supplemental Figure S4A). Details of the animals used in the SLAMseq experiment are given in Table 3.

**Figure 3.**
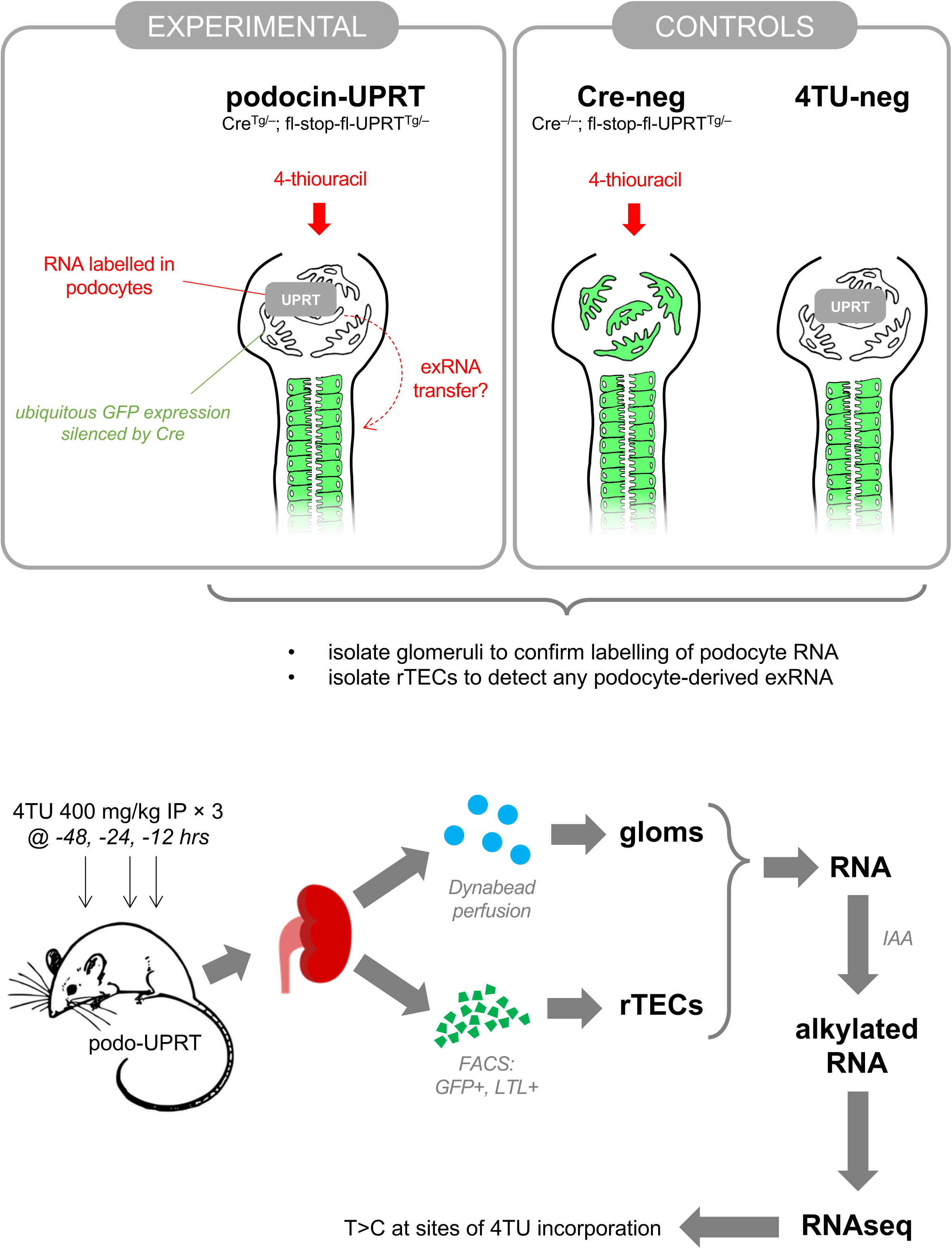
Experimental design for the labelling and detection of podocyte RNA using SLAMseq. Podocin-UPRT mice (and Cre-negative littermate controls) were treated with 4-thiouracil: 400 mg/kg body weight at 48, 24 and 12 hours prior to cull. A further control group received no 4-thiouracil. Kidneys were used either to provide glomeruli (isolated after Dynabead perfusion) or renal tubular epithelial cells (isolated by FACS). Glomeruli or rTECs were lysed and RNA extracted, alkylated and sequenced (SLAMseq).

**Table 3.**
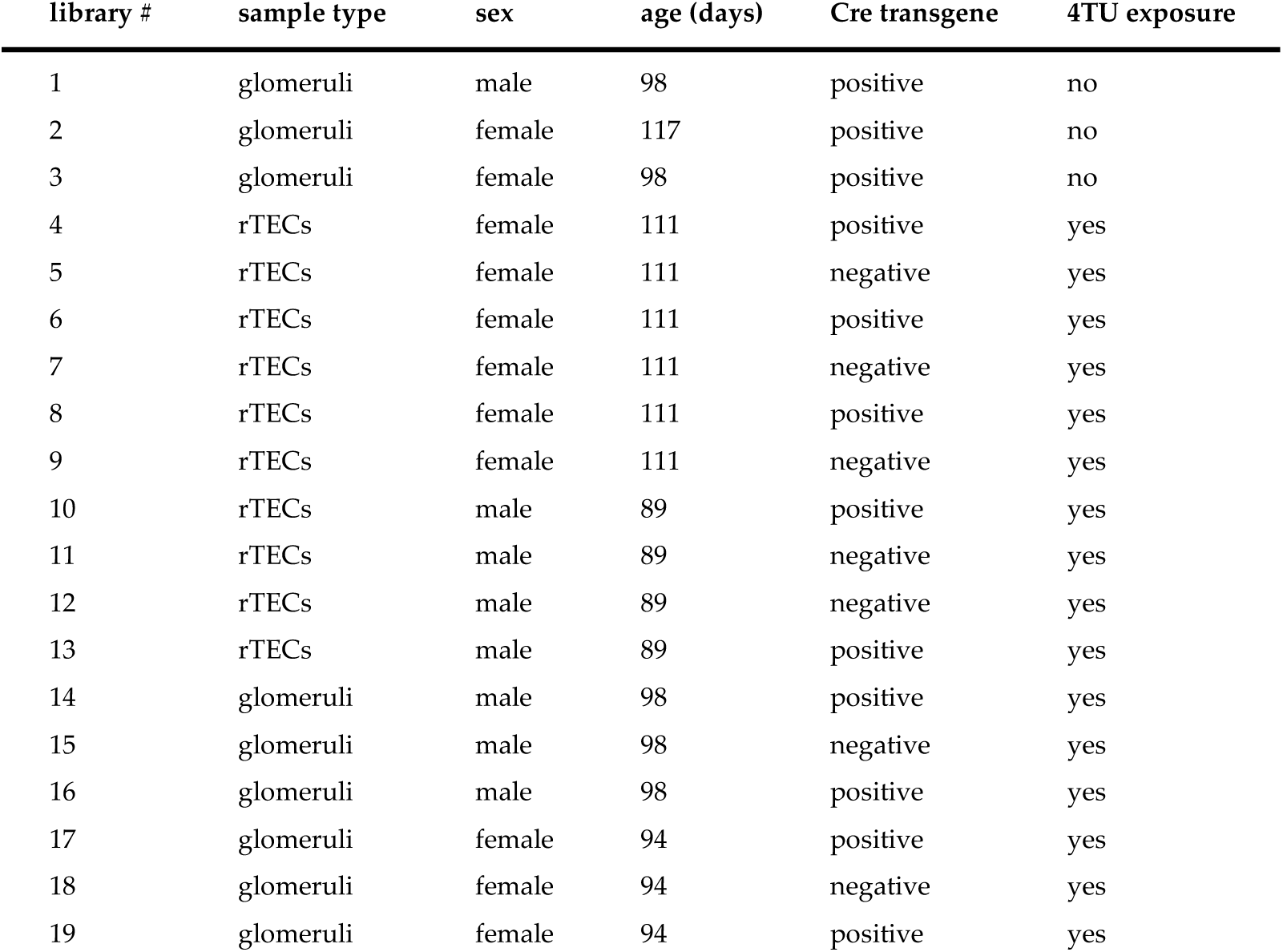
Mice used in SLAMseq experiment.

From these samples, total RNA was extracted and treated with iodoacetamide. The ability of the iodoacetamide treatment to induce alkylation of RNA was confirmed by using UV spectrophotometry to confirm alkylation of 4-thiouracil controls (supplemental Figure S4B). Alkylated mRNA was analysed by RNAseq (Quantseq); the SLAMDUNK analysis pipeline was used to quantify T>C conversions – expected at sites of 4TU incorporation – within 3’UTRs.^37^

Our RNAseq data confirmed that the transcriptomes of glomeruli and rTECs were distinct (supplemental Figure S4C) and enriched in known podocyte- and rTEC-restricted genes respectively (supplemental Figure S4D&E).

### 4-thiouracil successfully labels podocyte transcripts

In glomeruli, T>C conversions were observed with greater frequency in 4TU-treated podocyte–UPRT mice, compared to 4TU-treated Cre-negative mice and 4TU-naïve controls (Figure 4A,B&E). In a zero-inflation Poisson regression model, T>C conversion was increased both the Cre-negative and podo-UPRT groups, relative to the 4-TU naïve group. The zero-inflation model coefficients were –0.151 and –0.452 in the Cre-negative and podo-UPRT groups respectively (p < 2 × 10^−16^ for both); in other words, the odds of a gene exhibiting zero T>C conversions were reduced by 15% and 36% in Cre-negative and podo-UPRT mice respectively, compared to 4TU negative controls. The count model coefficients were 0.257 and 0.658 (p < 2 × 10^−16^ for both); in other words, in a model excluding any “excess” zero T>C conversions, the rate of T>C conversion was increased by 29% and 93% in Cre-negative and podo-UPRT mice respectively.

**Figure 4.**
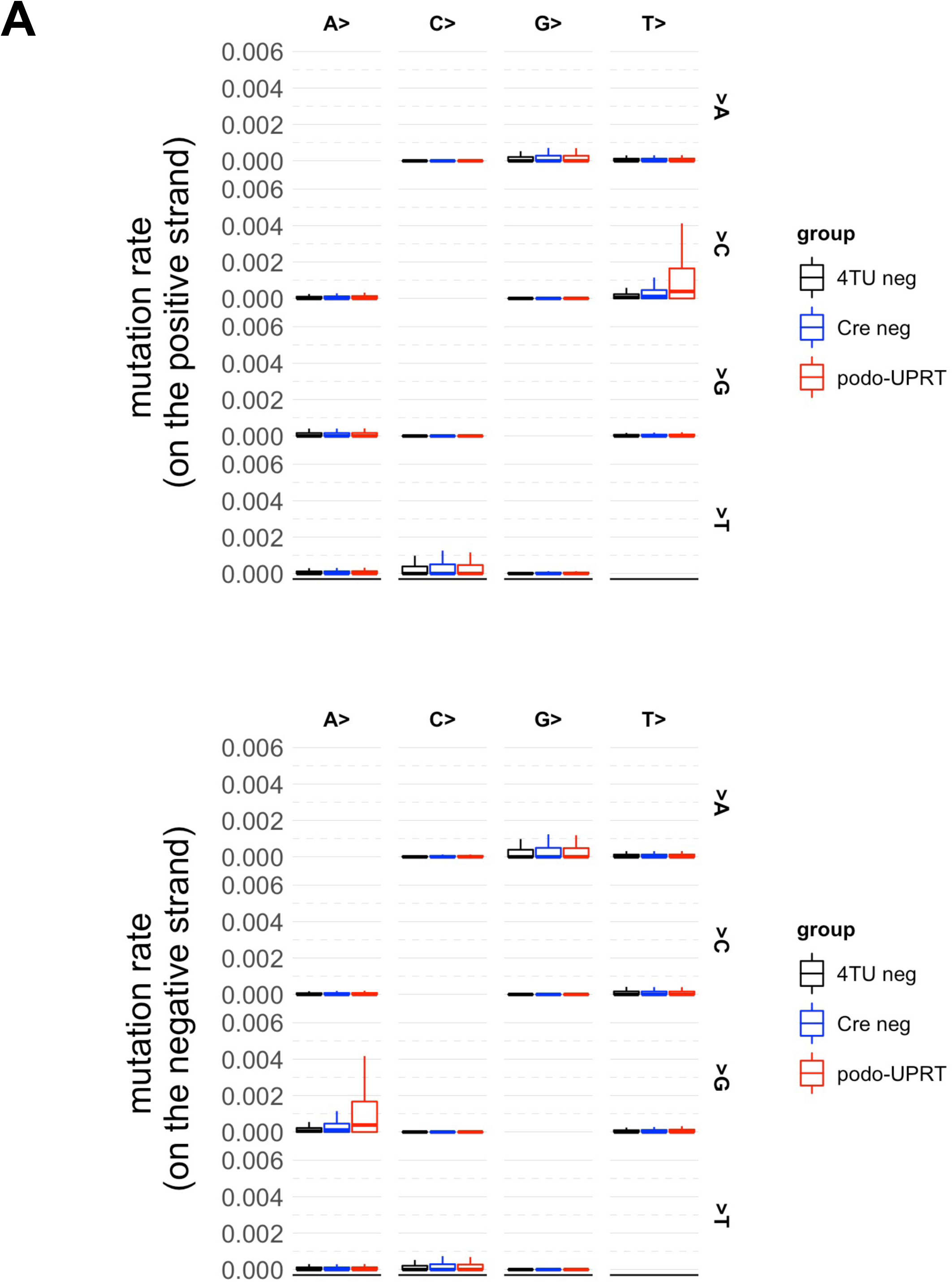
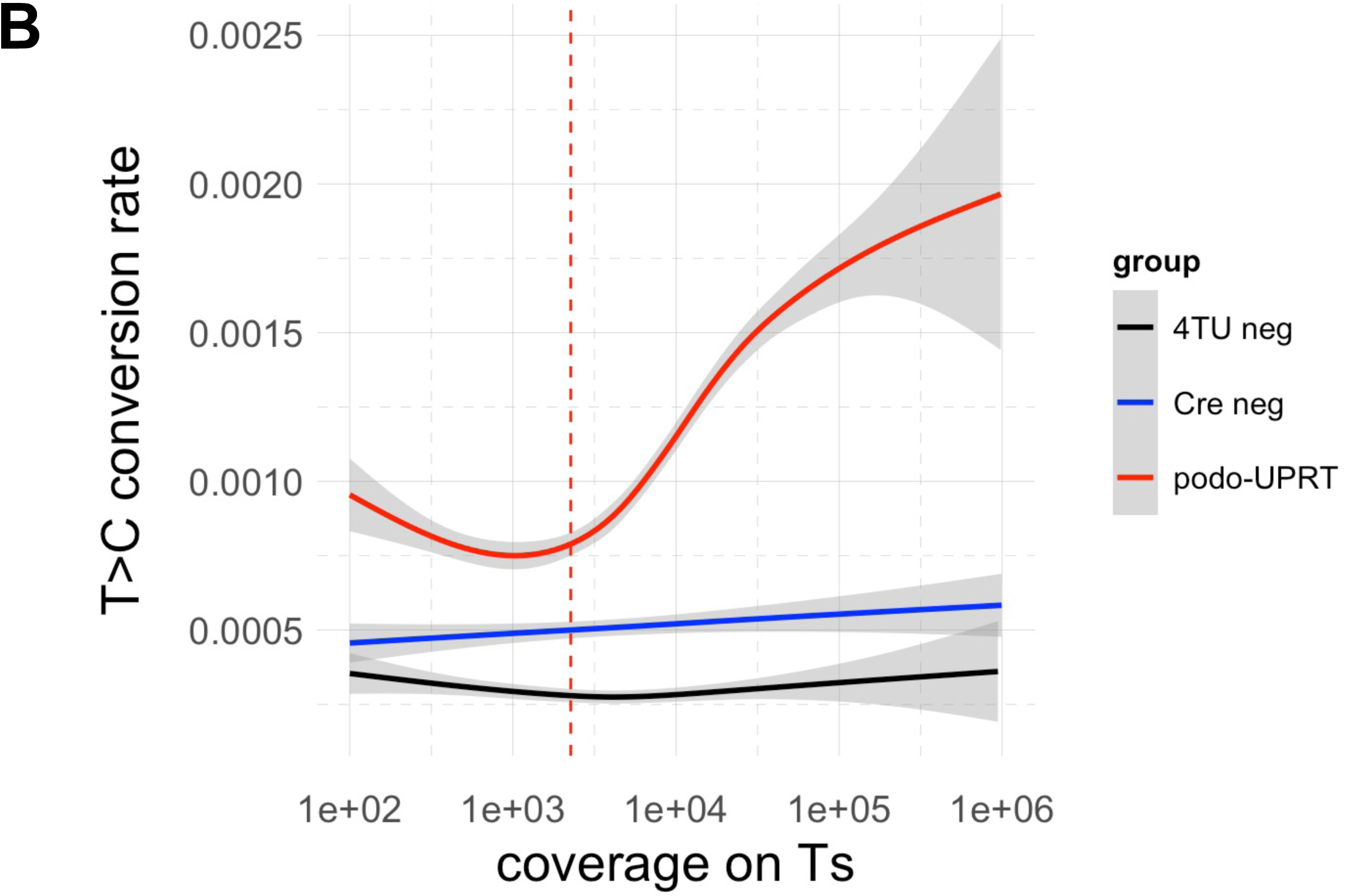
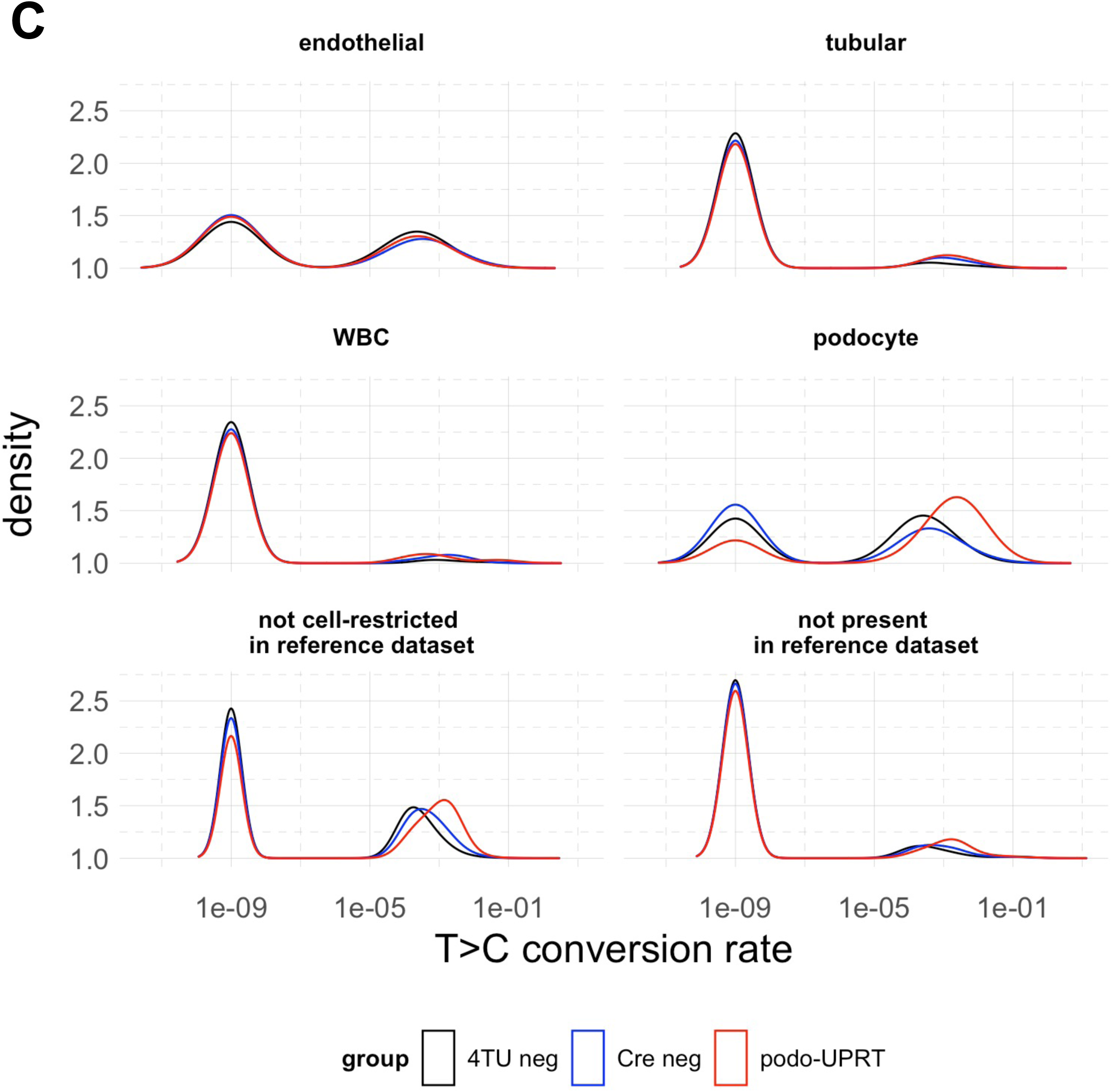
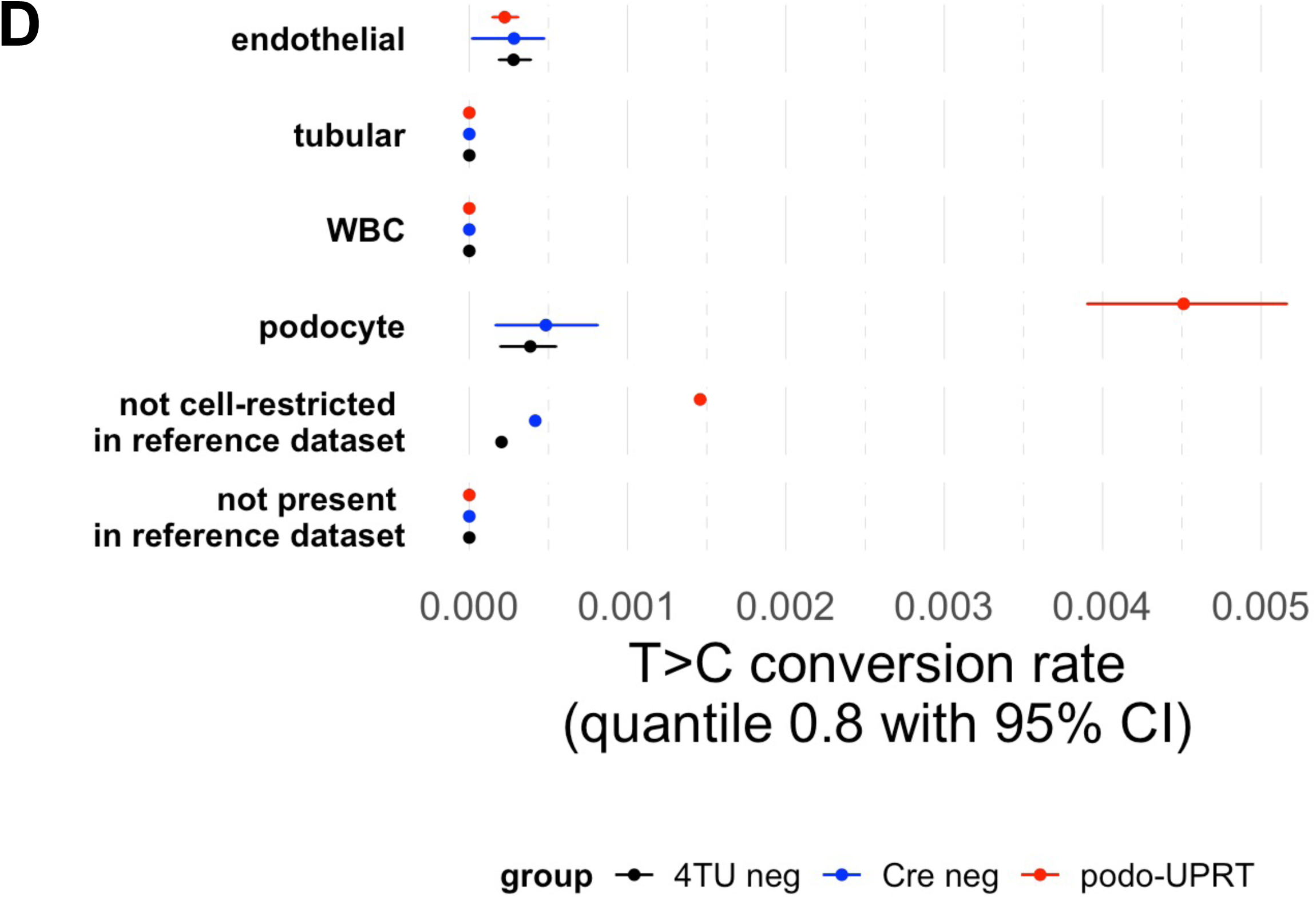
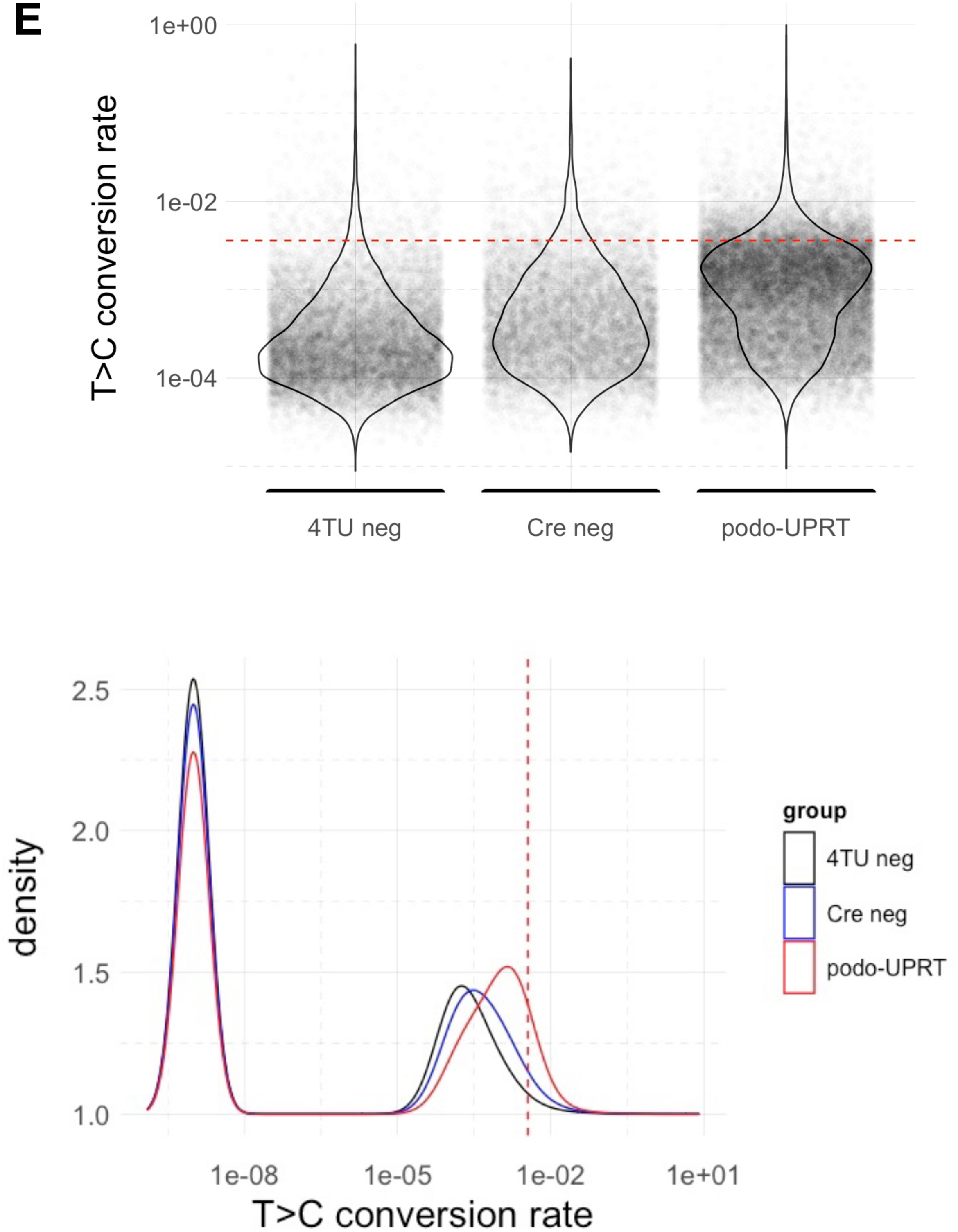
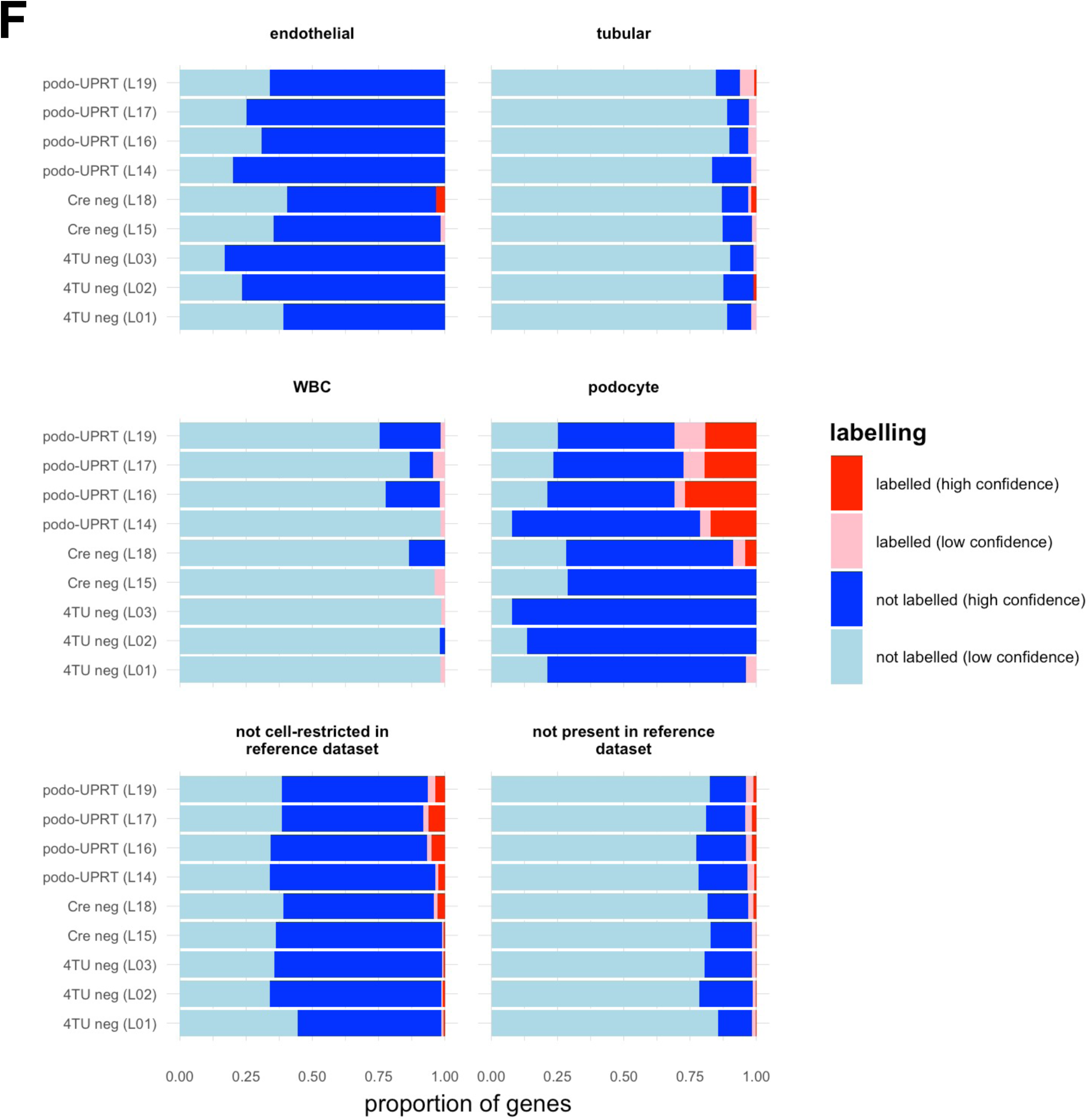
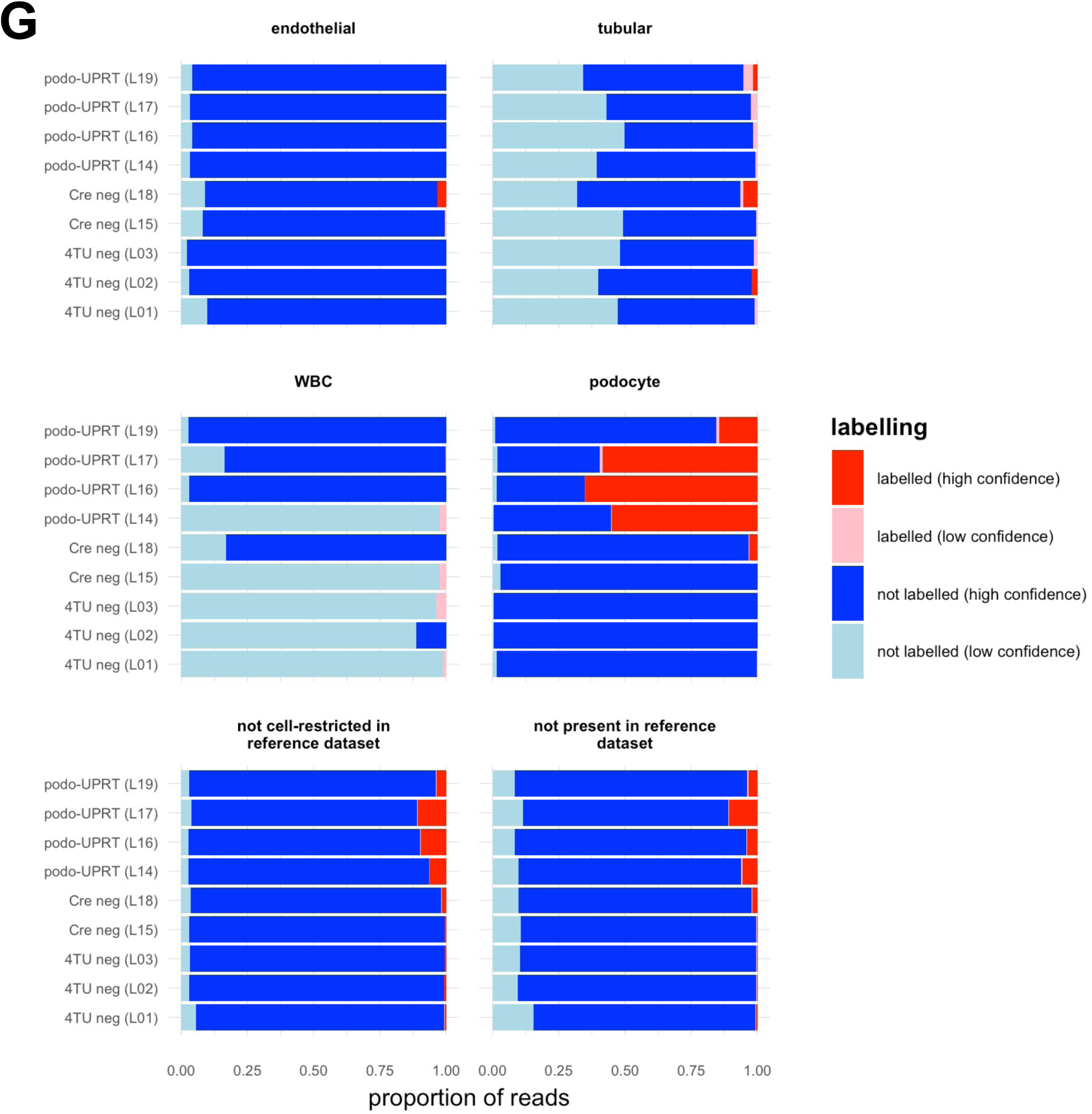
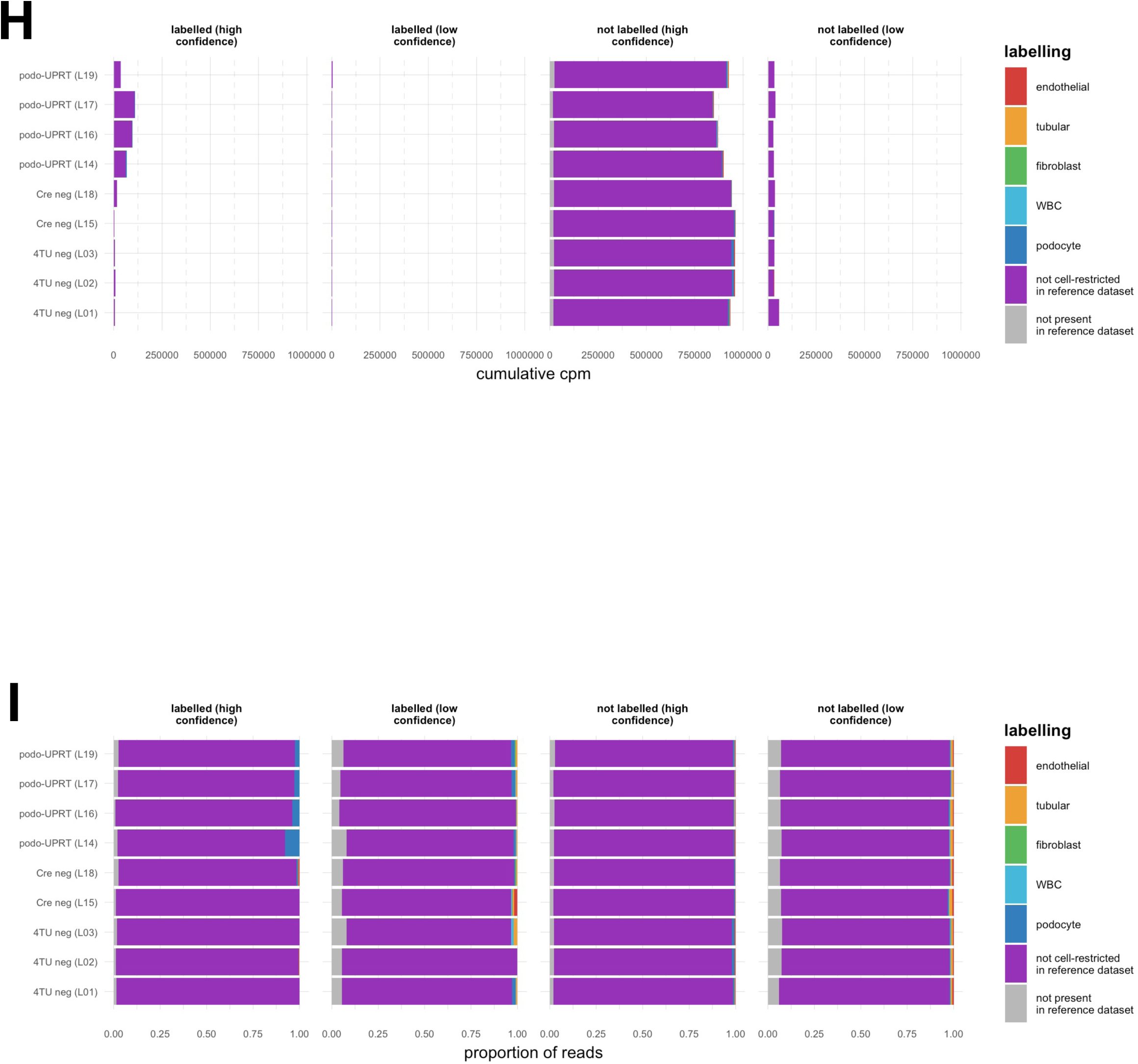
Labelling of known podocyte genes in glomerular samples. **A) Mutation rates – i.e. single nucleotide mismatches between our RNAseq data and the reference genome in mapped reads.** Podo-UPRT (Cre positive) libraries were enriched for T>C mutations on the positive strand (and A>G mutations on the negative strand), consistent with successful RNA labelling with 4-thiouracil. **B) Rates of T>C conversion in glomerular libraries.** Relationship between T>C conversion rate and the total number of T residues mapping to any given gene (“coverage on Ts”). The red, dashed vertical line indicates the inverse of the mean T>C conversion rate in 4TU-negative control libraries; in other words for this “coverage” we expect an average of one T>C conversion even in control libraries. Below this threshold, T>C conversion rates are sensitive to even a single T>C conversion and therefore RNA labelling cannot be inferred with high confidence. **C) T>C conversion rates in glomerular libraries in “marker genes”: distribution.** Cell-restricted “marker genes” were defined in a reference scRNAseq dataset. A constant correction factor (10^−9^) was added to each value before plotting, so that zero values appear on the log scale (large peak at 1 × 10^−9^). RNA labelling was evident in known podocyte genes, in genes expressed in multiple cell types and in genes that were not present in the reference dataset. Cre-dependent RNA labelling was not evident in endothelial, fibroblast, tubular or WBC marker genes. **D) T>C conversion rates in glomerular libraries in “marker genes”: summary statistic.** Due to the non-normal distribution of T>C conversion rates and a high proportion of reads with zero T>C conversions, we selected quantile 0.8 as a summary statistic as this lay approximately in the middle of the right-hand (non-zero) peak in the bimodal distribution. **E) Method for defining “labelled” genes.** We defined genes as being confidently labelled if they exhibited T>C conversion at a rate exceeding the 99^th^ centile for genes in 4TU-naïve control libraries. This threshold is represented by the dashed red line. (In the lower plot, a constant correction factor was added before plotting, as in C). **F) & G) Proportion of genes (F) or reads (G) exhibiting labelling.** We determined the proportion of genes exhibiting labelling, stratifying the results by genes known to be cell-enriched in a reference dataset. As in C), labelling was evident in podocyte marker genes and ubiquitously-expressed genes but not in markers for other cell types. **H & I) Proportion of reads exhibiting labelling, stratified by labelling status.** The same data in G are re-presented, this time using colour to represent known “marker” genes and the plot facet to represent labelling status; only a minority of total reads exhibit labelling with high confidence.

In order to verify that our approach labels known podocyte-restricted / podocyte-enriched genes, we used an independent reference scRNAseq dataset^40^ to define a set of “marker” genes for various kidney cell types. We detected increased rates of T>C conversion in known podocyte-restricted genes but not in genes known to be restricted to other cell types (Figure 4C&D; p-values for Kruskal-Wallis test within each marker gene set: endothelial cells: 0.57; fibroblasts: 0.62; podocytes: <10^-23^; tubular cells: 0.004; white blood cells, 0.02; genes expressed in multiple cell types: <10^-23^; genes not expressed in the reference dataset: <10^-14^).

The rate of T>C conversion varied with the total number of T residues sampled within the 3’UTR of a given gene: the “coverage on Ts”. When very few T residues were sampled, the conversion rate was very sensitive to even a single T>C conversion. Therefore, within each library, we classified genes according to whether we could call labelling with high confidence (high coverage on Ts) or low confidence (low coverage on Ts). We set this threshold as the reciprocal of the mean conversion rate in the 4TU-negative group – i.e. that is the coverage at which an average of one T>C conversion would be observed at background rates (red dashed line in Figure 4B). Unsurprisingly, a low-level of “background” T>C conversion was observed even in negative controls. Therefore, we took a conservative approach to defining “labelled” genes. Within the “high confidence” genes in each library, we classified a gene as being labelled if the rate of T>C conversion exceeded the 99^th^ centile for all “high confidence” genes in 4TU-naïve control libraries (Figure 4E). This approach effectively sets a false discovery rate of 1% for identifying labelled genes.

In glomerular libraries, the proportion of genes deemed to be labelled with high confidence was 7.1% (95% confidence interval 6.8 – 7.4%) in 4TU-treated podocyte-UPRT mice, 2.5% (2.3 – 2.7%) in Cre-negative controls and 1.0% (0.9 – 1.1%) in 4TU-naïve controls. Labelled genes corresponded to known podocyte-enriched genes, genes known to be expressed in multiple cell types or genes that were not present in the reference dataset (Figure 4F&G). The vast majority of genes exhibiting labelling were expressed in multiple cell types in the reference dataset (Figure 4H&I).

### Labelled RNA is detected in renal tubular cells

Cre expression removes a floxed GFP-triple-stop cassette from the floxed-stop-UPRT transgene. Therefore, we isolated rTECs using a two-fluorophore FACS approach, selecting LTL-positive, GFP-positive cells (supplemental Figure S3). This stringent approach minimised any possibility of including Cre-positive (and therefore GFP-negative) podocytes in the rTEC sample. As expected, the proportion of GFP-positive cells in whole kidney digests was lower in podocyte-UPRT mice than in Cre-negative littermates (Figure 5A; p = 0.037 for effect of Cre, p = 0.951 for effect of LTL-positivity, p = 0.953 for interaction by 2-way ANOVA). This difference was confined to LTL-negative cells.

**Figure 5.**
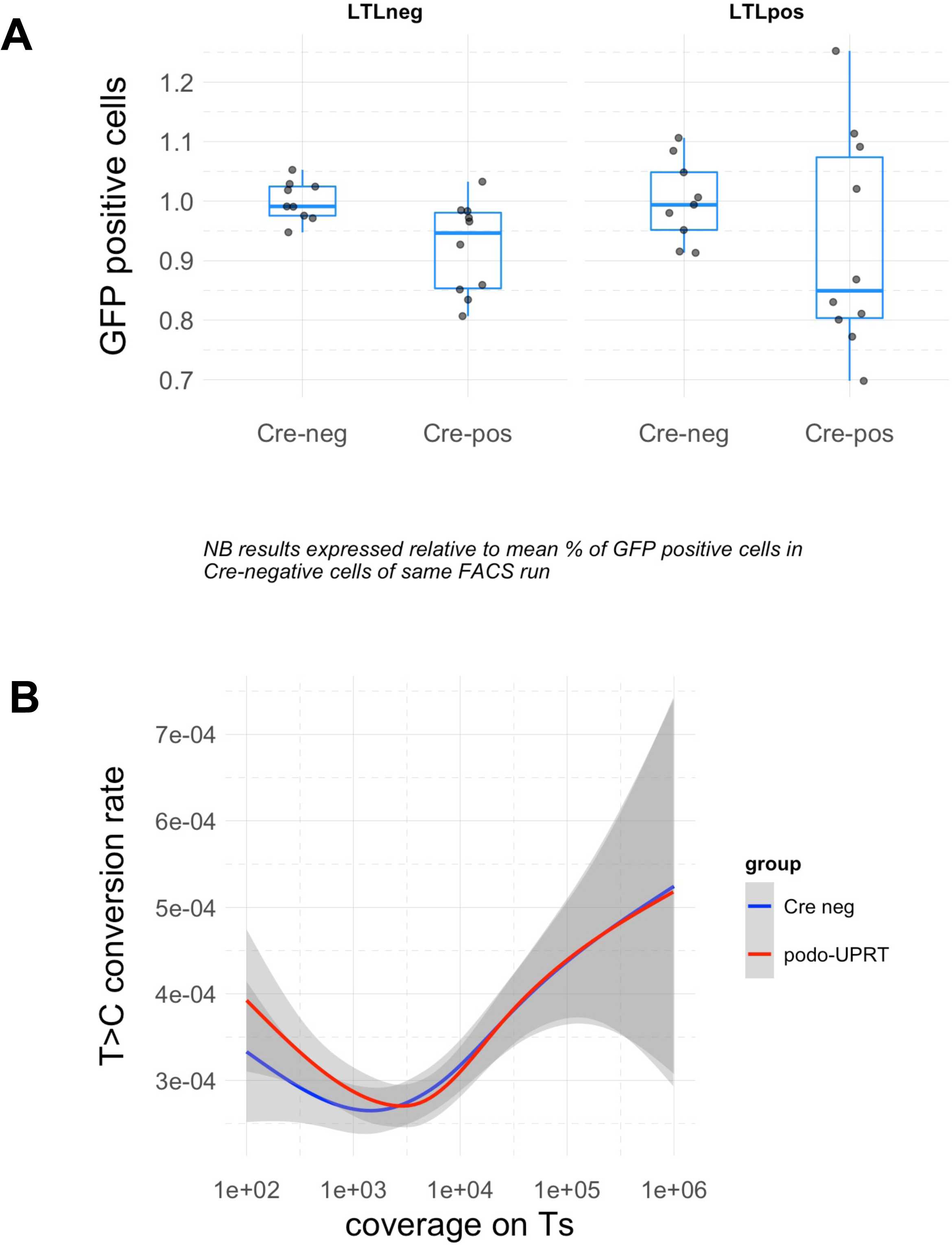
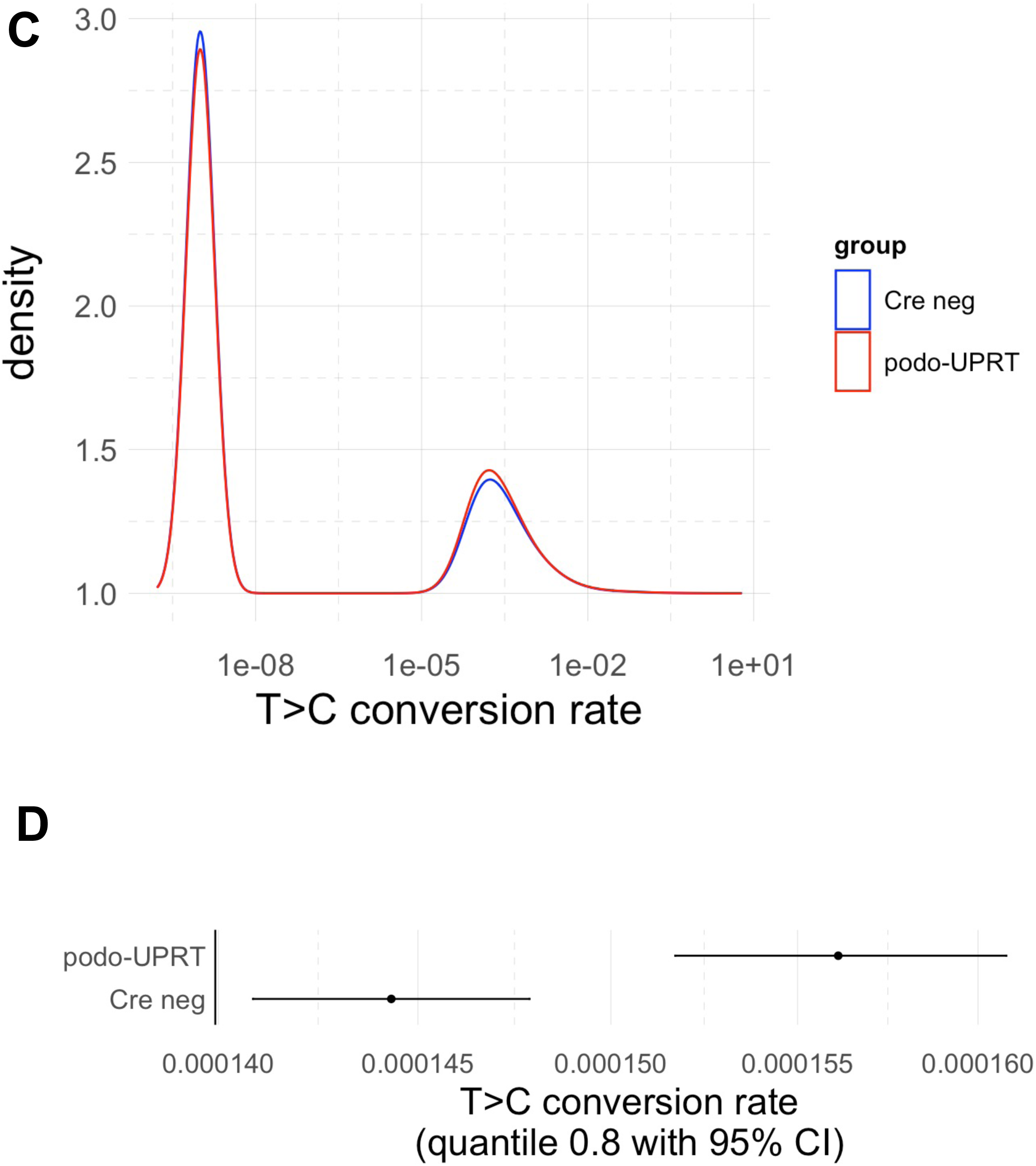
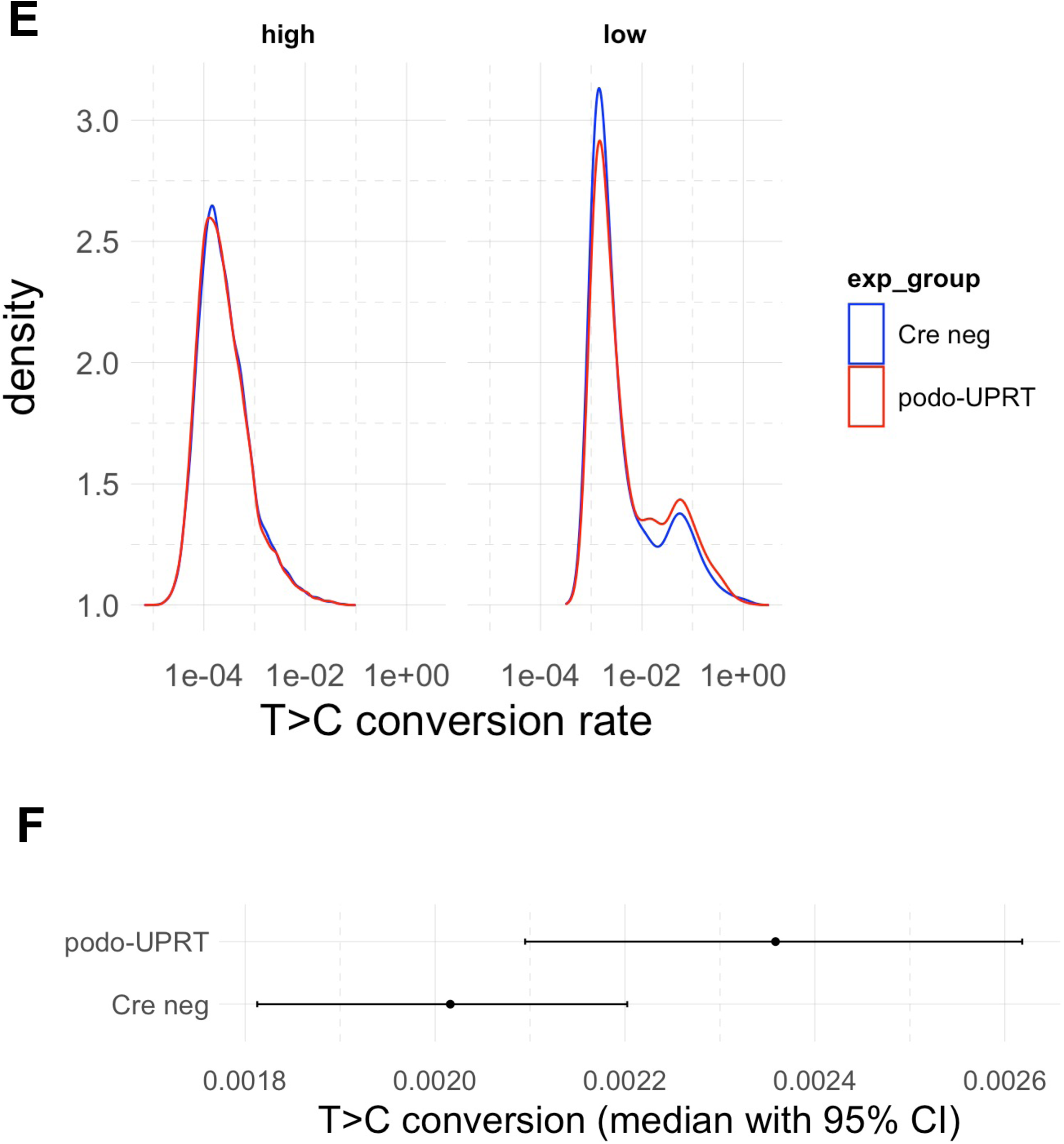
Isolation of rTECs by FACS. **A) Proportion of GFP+ve cells, stratified by genotype and LTL-positivity.** Cre-mediated loss of GFP expression was evident in the LTL-negative (i.e. mixed origin) cells but not in LTL-positive (i.e. rTEC) cells. **B) Global rates of T>C conversion in rTECs.** A small excess of T>C conversion was evidence in reads with low T coverage. **C) Global rates of T>C conversion in rTECs: distribution.** A constant correction factor (10^−9^) was added to each value before plotting, so that zero values appear on the log scale (large peak at 1 × 10^−9^). A small shift from completely unlabelled reads (the tall spike at the left) to reads with low-level T>C conversion (the wider hump to the right) is evident in podo-UPRT libraries. **D) Global rates of T>C conversion in rTECs: summary statistic.** T>C conversion rates (80^th^ centile with 95% confidence interval) were significantly higher in podo-UPRT mice than in Cre-negative controls but the absolute increase was very small. The T>C conversion (80^th^ centile) corresponded to labelling of ∼1 in 6900 uridines in Cre negative controls and ∼1 in 6400 uridines in podo-UPRT mice. **E) T>C conversion rates in tubular libraries, stratified by T coverage.** We set an arbitrary threshold of 1000 Ts to define low vs. high coverage. **F) Median T>C conversion rates** within low-coverage tubular genes with at least one T>C conversion.

In rTECs, rates of T>C conversion were slightly higher in 4TU-treated podocyte-UPRT mice than in 4TU-treated Cre-negative controls (Figure 5B). The distribution of T>C conversion rates in rTEC libraries exhibited a small, Cre-dependent shift from genes with no T>C conversions to genes with a low rate of T>C conversion (Figure 5C&D). In a zero-inflated Poisson regression model, the zero-inflation model coefficient was –0.104 in the podo-UPRT group (p = 7.4 × 10^−16^); in other words, the odds of a gene exhibiting zero T>C conversions were reduced by 10% compared to Cre-negative controls.

We noted that higher rates of T>C conversion were noted for genes with low coverage on Ts (Figure 5B). We therefore examined those genes in greater detail, arbitrarily defining low coverage as any gene with fewer than 1000 sampled T residues. (We normalised to T coverage rather than read counts or counts-per-million so as to make a comparison of T>C conversion rates between experimental groups that was not biased by library size.) In this subset of low-coverage genes, there was a higher rate of T>C conversion in the podo-UPRT compared to the Cre-negative group (Figure 5E&F). The rate of T>C conversion was increased by 12.8% in the podo-UPRT group (p < 2 × 10^−16^ in a Poisson model excluding excess zeros).

These observations suggest that 4TU-labelled RNA moves from podocytes to rTECs. We attempted to determine the RNA sequences that were transferred into tubular cells. Looking within the set of low-coverage tubular reads, we excluded genes that had zero T>C conversions in all libraries and then defined a putative podocyte-derived gene if the mean rate of T>C conversion was at least 1.5x higher in podo-UPRT mice than in podo-Cre mice and if the number of podo-UPRT libraries with 1 or more T>C conversions exceeded the number of Cre-negative libraries with 1 or more T>C conversions by at least 2. 1011 genes were identified as being putative podocyte-derived messenger RNA (supplemental table S2). However, the confidence with which we could identify any single gene as being podocyte-derived was low: when we used the same criteria to identify genes that were apparently labelled in Cre-negative (but not podo-UPRT) libraries, we identified 666 genes.

## Discussion

### Main findings

We used RNA labelling to directly track podocyte-derived RNA in the mouse kidney. We detected labelled RNA in renal tubular epithelial cells, compatible with the *in vivo* transfer of RNA from podocyte to rTEC. This provides proof-of-principle that SLAMseq can be used to track RNA mobility *in vivo*, building on an earlier study in which this approach was used to show transfer of RNA from epididymis to sperm in the mouse.^12^ This is an important finding, providing the first direct evidence of physiological exRNA transfer between kidney cells. It provokes the obvious question: does such transfer induce important functional changes in the recipient renal tubular epithelial cell? This is plausible, given that single microRNAs delivered in extracellular vesicles can induce profound changes in rTEC phenotype.^1,2^

### The podo-UPRT mouse is a useful tool for studying glomerular disease

We generated the podo-UPRT mouse, in which podocyte RNA is specifically labelled with 4-thiouracil. We validated this approach by showing that podocyte-restricted genes (as defined in an independent reference dataset) were labelled whereas genes known to be restricted to other cell types were not.

This is a potentially powerful tool for studying the molecular mechanisms of glomerular disease. We demonstrated that despite having a predominantly C57BL/6 background (classically resistant to glomerular injury), podo-UPRT mice developed albuminuria with doxorubicin treatment. Therefore, these mice could be used to study toxic injury models, even without cross-breeding onto a more susceptible genetic background.

The podo-UPRT mouse offers advantages over alternative methods of transcriptional profiling. It does not rely on FACS-sorting or laser capture microdissection of podocytes and could – in theory – be used to profile all classes of RNA (not just mRNA, as is profiled in TRAP for example).^52^

### Limitations

There are a number of important limitations in our approach to studying RNA transfer between kidney cells. First, we have so far restricted our analysis to messenger RNA; it will be important to investigate transfer of other RNA classes in the podo-UPRT mouse, particularly the small, non-coding RNAs that are enriched in the extracellular fraction. Second, our approach was unable to differentiate between labelled exRNA that was firmly adherent to the surface of rTECs (rather than labelled RNA that had been truly internalised by rTECs), although it seems likely that any cell-extrinsic RNA would be washed away during preparation for FACS.

Third, our bulk-sequencing approach was unable to determine the heterogeneity of exRNA uptake by rTECs. We chose a stringent double-positive (LTL+, GFP+) protocol to isolate rTECs by FACS in order to minimise any possibility of co-purifying UPRT-expressing podocytes. It is possible that this approach led us to *underestimate* the extent of any RNA transfer if rTECs that had taken up large quantities of podocyte-derived RNA also took up Cre mRNA, with subsequent loss of GFP expression.

Finally, the signal-to-noise ratio in tubular SLAMseq data was high. Taking the 80^th^ centile as a summary statistic, rates of T>C conversion was ∼1 in 6900 in Cre-negative control libraries, rising to ∼1 in 6400 in podo-URPT mice. Therefore, although we were able to find evidence of RNA transfer across our entire dataset, we were unable to confidently determine *which* RNA sequences had been transferred from podocyte to renal tubular cell. (This limitation was also encountered in the attempt to use SLAMseq to track RNA moving from epididymis to sperm.^12^) We detected increased T>C conversion rates within tubular reads with low T coverage (broadly equating to low-abundance genes); this is compatible with a model in which unlabelled rTEC-endogenous reads “swamp” any signal from labelled RNA in genes expressed at high abundance in rTECs. The signal-to-noise ratio could, in theory, be improved by sequencing to greater depth or by modifying the rTEC isolation protocol to enrich for cells that have taken up large quantities of podocyte-derived RNA.

### Perspectives

We have provided important evidence that RNA moves from podocyte to renal tubule. Our results raise a number of tantalising questions: how much RNA is taken up by rTECs?; which RNA sequences move?; how heterogeneous is RNA uptake?; what is the route of exRNA transfer? It is likely that exRNA is shuttled between cells in extracellular vesicles, lipoprotein particles or bound to RNA-binding proteins.^53^ The most direct route of transfer from podocyte to rTEC is through the urinary space but we cannot discount the possibility that exRNA moved through the vascular space. (Extracellular vesicles have an apparent ability to traverse the glomerular and tubular basement membranes, despite the fact that this ought to be impossible given their size and charge.^19,54^)

The critical question is whether exRNA transfer mediates any meaningful biological function in homeostasis or disease. Whilst our data do not address this question, they are an important first piece in the puzzle. Teleologically, it would make sense for rTECs to respond to injury signals encoded in exRNA by activating pro-inflammatory, pro-repair pathways. Such a response would be maladaptive in the face of persistent podocyte injury, potentially contributing to the tubulointerstitial pathology that is observed in chronic glomerular disease.

## Conclusion

We have achieved podocyte-specific RNA labelling by using SLAMseq in the podo-UPRT mouse. We detected labelled RNA in renal tubular cells. This suggests that RNA moves from podocyte to renal tubular epithelial cells in the mammalian kidney. Our model could be used to explore the functional consequences of this novel mode of cell-to-cell communication within the kidney.

## Acknowledgements & funding

This work was funded by a Wellcome Clinical Research Career Development Fellowship awarded to RWH (209562/Z/17/Z).

We are very grateful for the technical assistance provided by the University of Edinburgh Shared University Research Facilities (SuRF): Lyndsey Boswell, Fiona Inglis & Michael Millar in Histology (for assistance with immunofluorescence and fluorescence microscopy), Pamela Brown in the Biomolecular facility (RNA capillary electrophoresis), Mari Pattison, Will Ramsay & Shonna Johnstone in the FACS facility and Forbes Howie in the Specialist Assay Service (creatinine and albumin assays). We are grateful to the technical assistance provided by the Bioresearch & Veterinary services at University of Edinburgh for their help in the design and implementation of mouse experiments and for re-deriving podocin-Cre mice from cryopreserved sperm.

## Author contributions

**RWH:** conceptualization, methodology, software, validation, formal analysis, investigation, writing – original draft, visualisation, funding acquisition; **SK:** software, visualisation; **RJMC:** conceptualisation, resources, supervision, writing – review & editing; **AB:** conceptualisation, resources, supervision, writing – review & editing; **JWD:** conceptualisation, methodology, resources, supervision, writing – review & editing.

## Supplemental material

**Table S1 – Genetic strain analysis of podocyte-UPRT mice. Report from Transnetyx SNP panel.**

**Table S2 – List of putative podocyte-derived genes in tubular libraries.**

---

**Figure S1.**
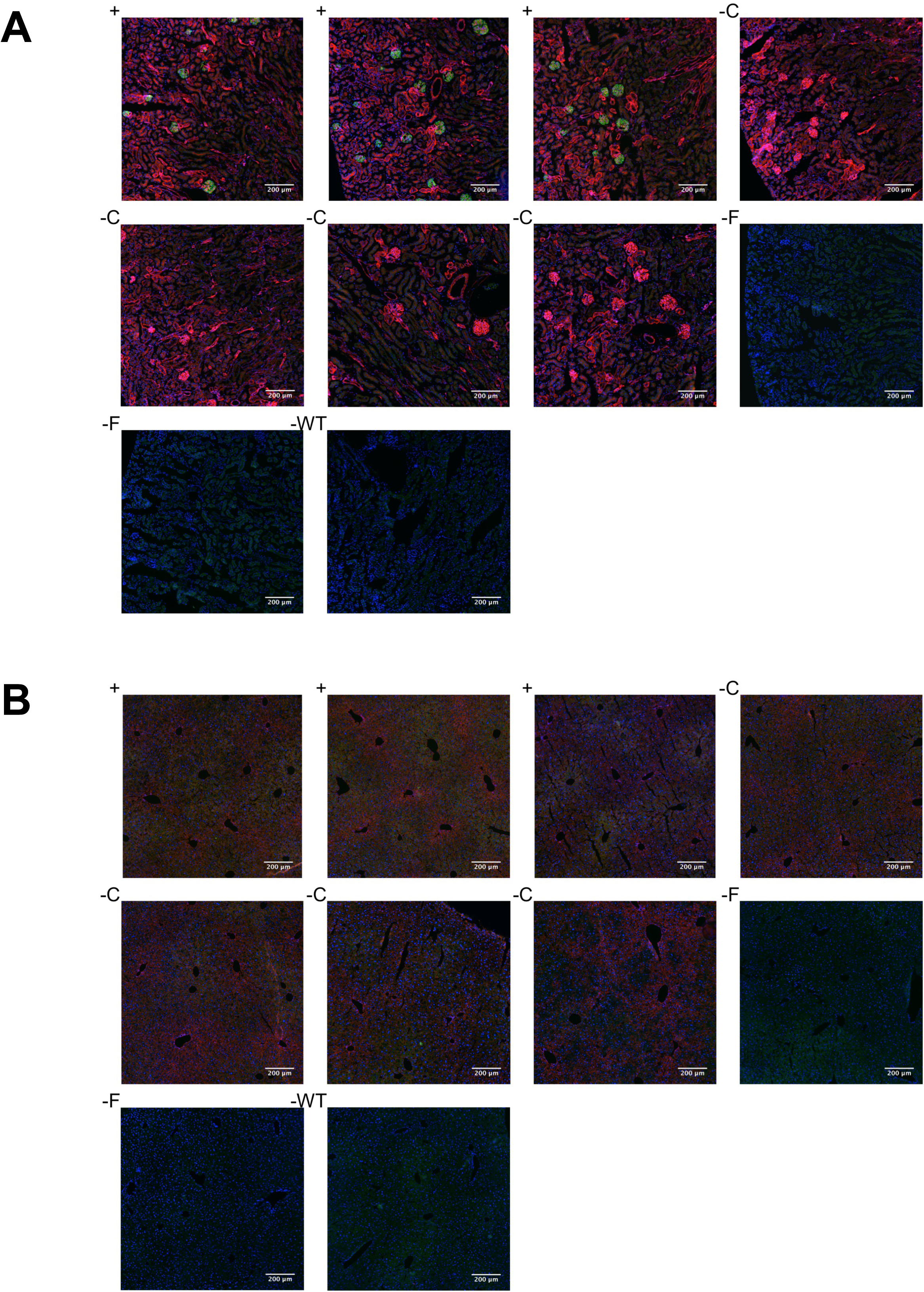
Cre expression in mTmG mice. **A) Kidney sections.** Cre-dependent green fluorescence was observed only in podocytes. **B) Liver sections.** There was no off-target Cre-dependent green fluorescence. Genotype codes are: “+” = Cre^Tg/−^, mTmG ^Tg/−^; “–C” = Cre^−/−^, mTmG ^Tg/−^; “-F” = Cre^Tg/−^, mTmG ^−/−^; WT = wild-type.

**Figure S2.**
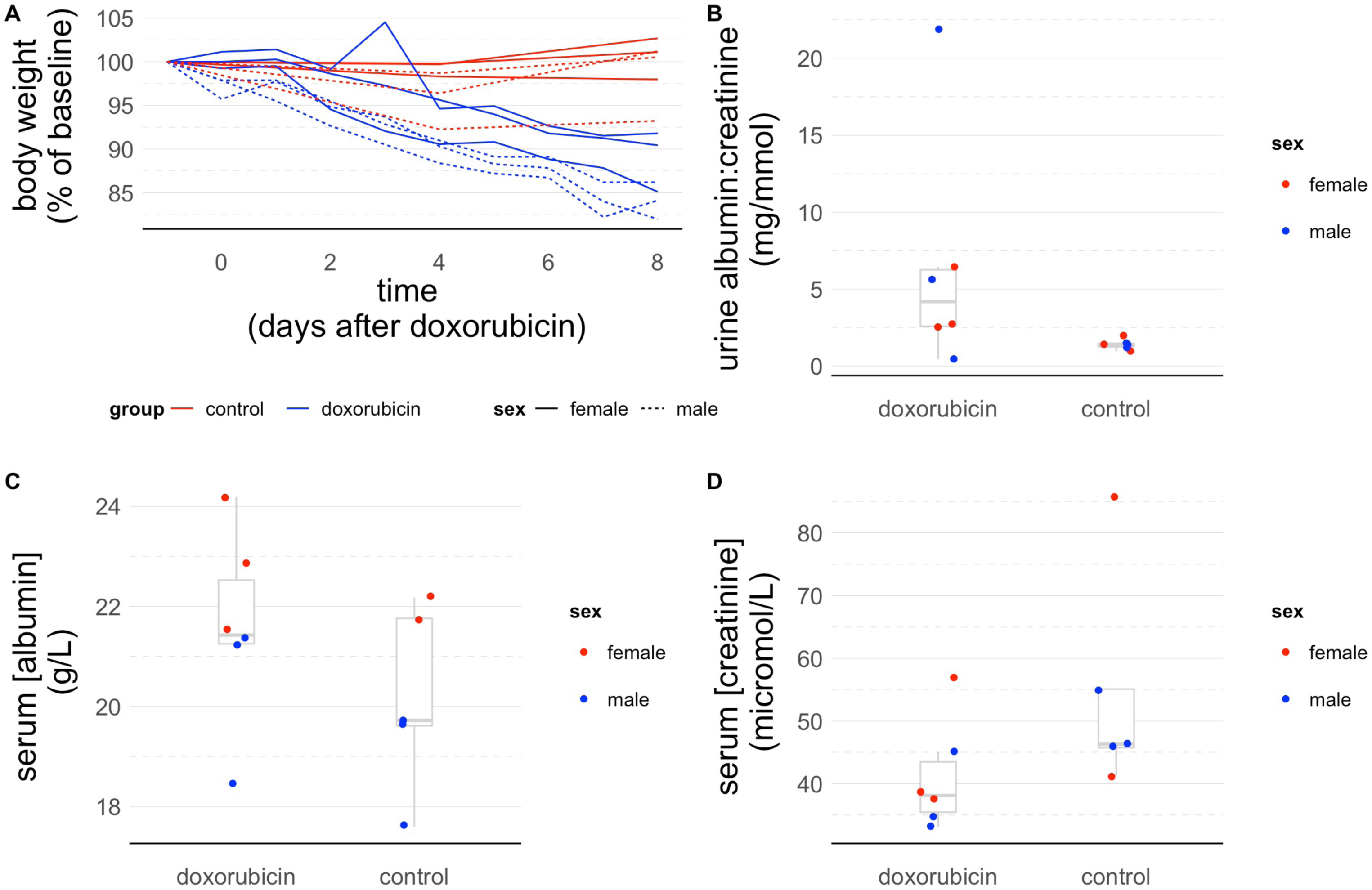
Doxorubicin injury model. Mice were treated with a single intravenous injection of doxorubicin (adriamycin) at 15 mcg per g body weight. They were monitored for 8 days before euthanasia, at which time a terminal blood sample was obtained. **A)** Body weight. **B)** Urine albumin excretion. **C)** Terminal serum creatinine and albumin concentrations.

**Figure S3.**
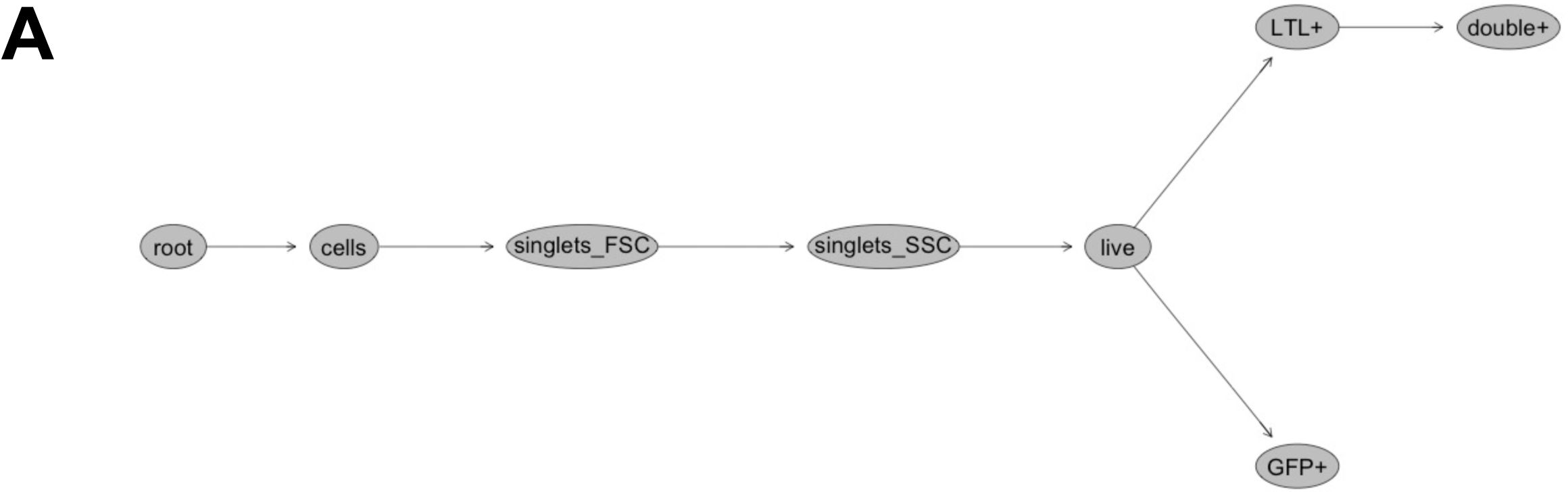
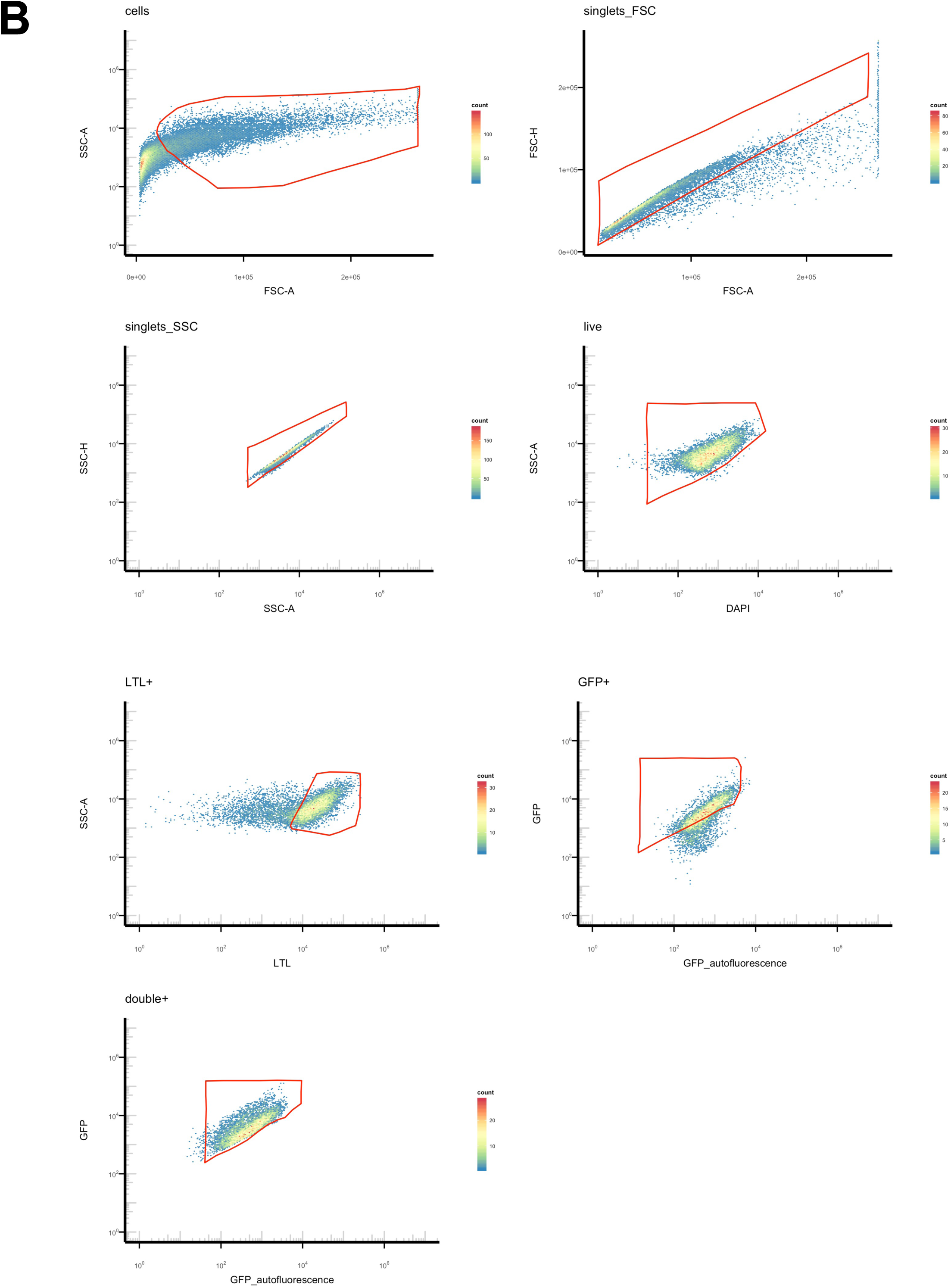
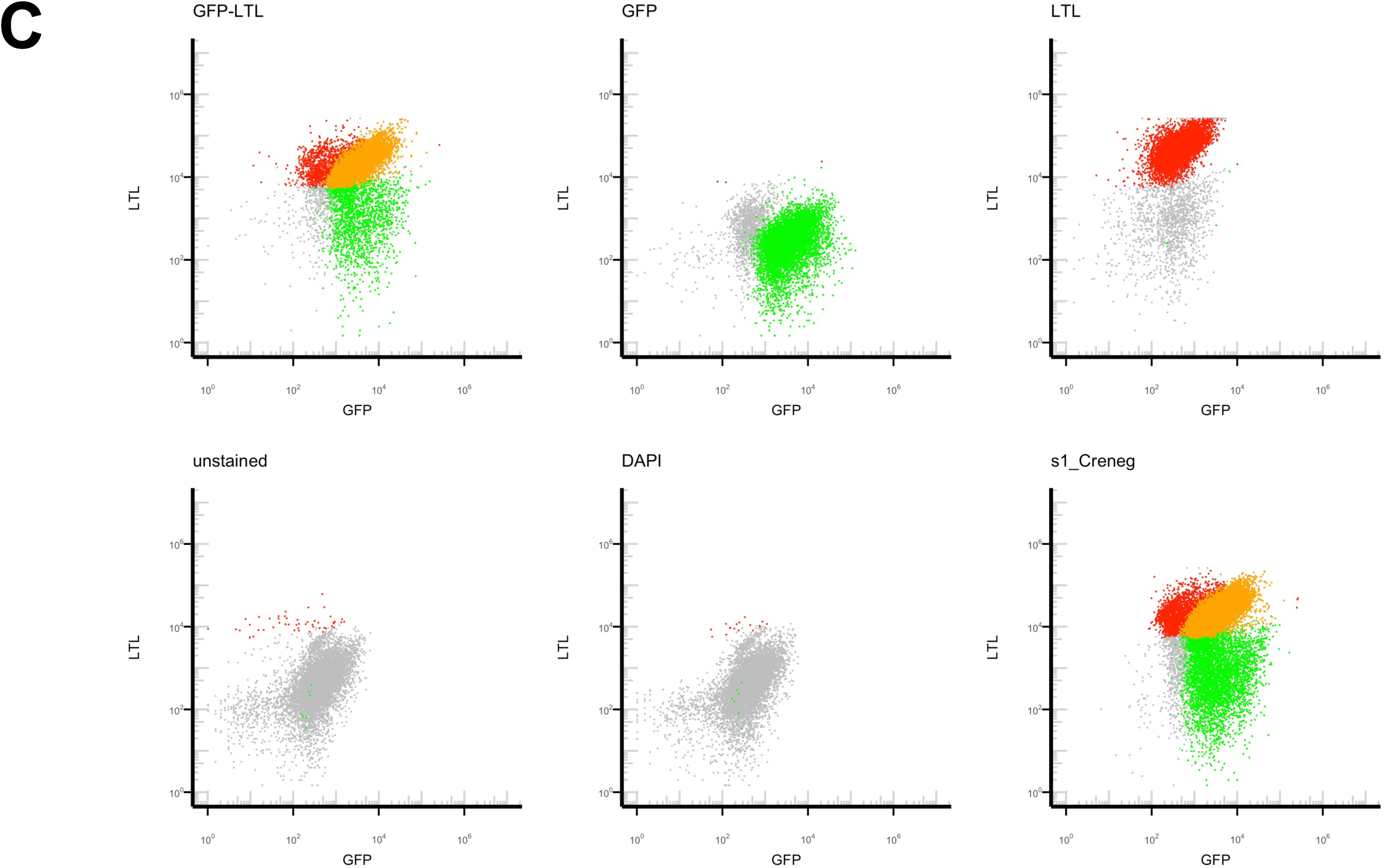
FACS strategy. All cells will constitutively express GFP, unless Cre recombinase has been active. We therefore selected rTECs by their ability to bind Lotus Tetragonolobus Lectin and express GFP. **A) Gating strategy. B) Representative gates. C) Control and experimental samples.** Cells classed as LTL+ve (red), GFP+ve (green), double-positive (orange) or double-negative (grey). Control samples were: “GFP-LTL” = stained with LTL and expressing GFP (i.e. as for experimental samples); “GFP” = GFP only (i.e. LTL-negative); “LTL” = LTL only (i.e. LTL staining in a GFP-negative mouse); “unstained” (i.e. negative for both LTL and GFP); “DAPI” = unstained + DAPI. Sample “s1_Creneg” was one of the experimental samples.

**Figure S4.**
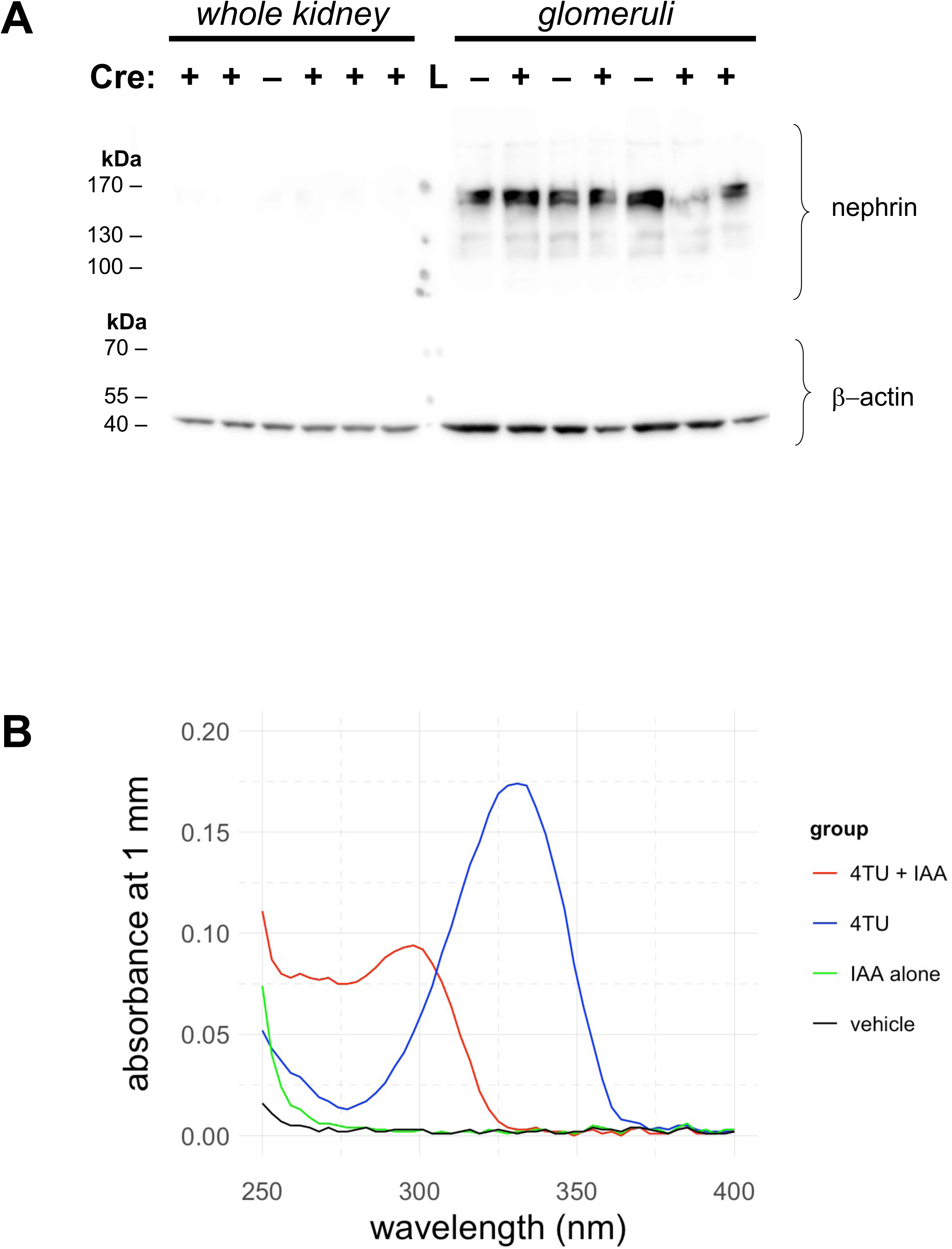
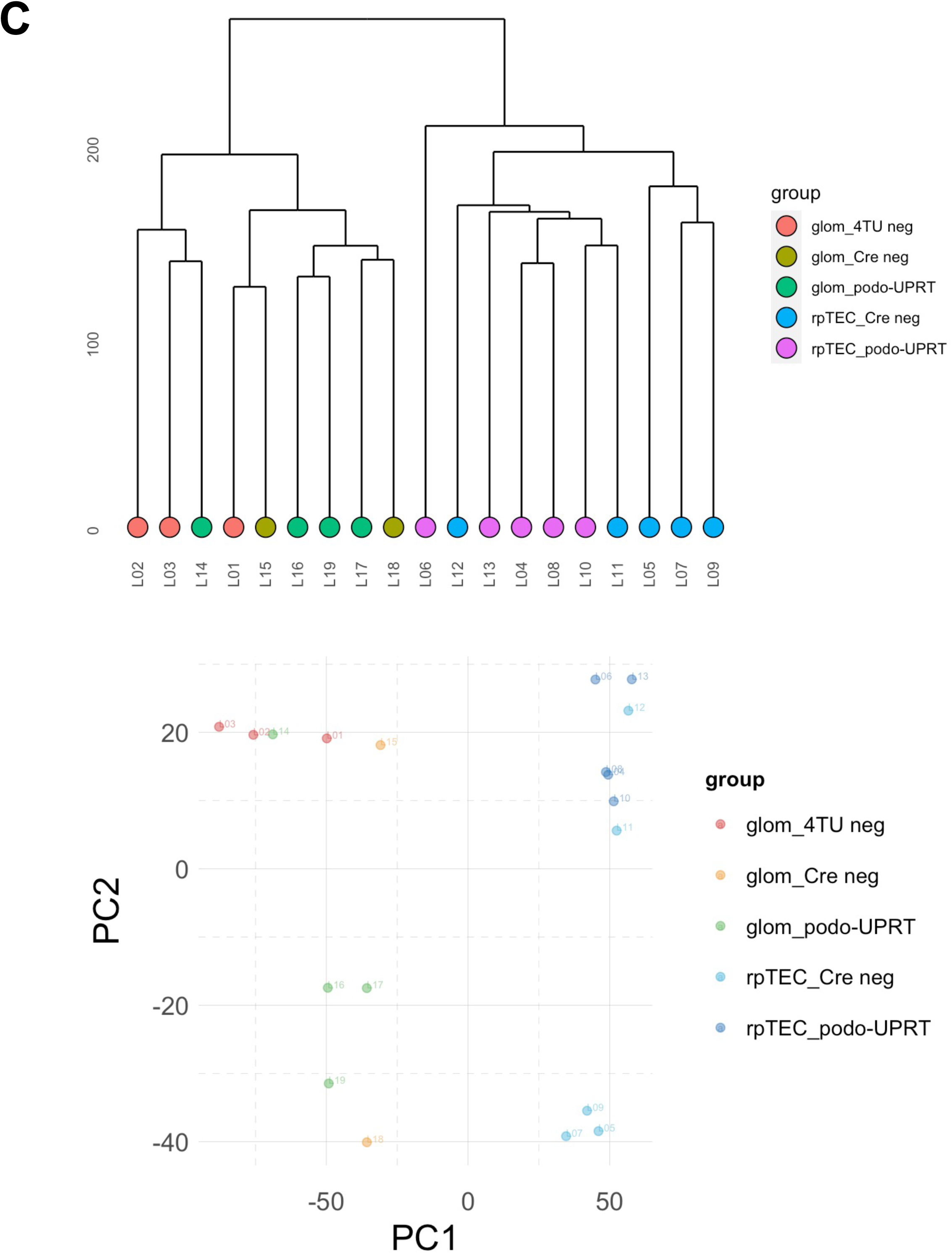
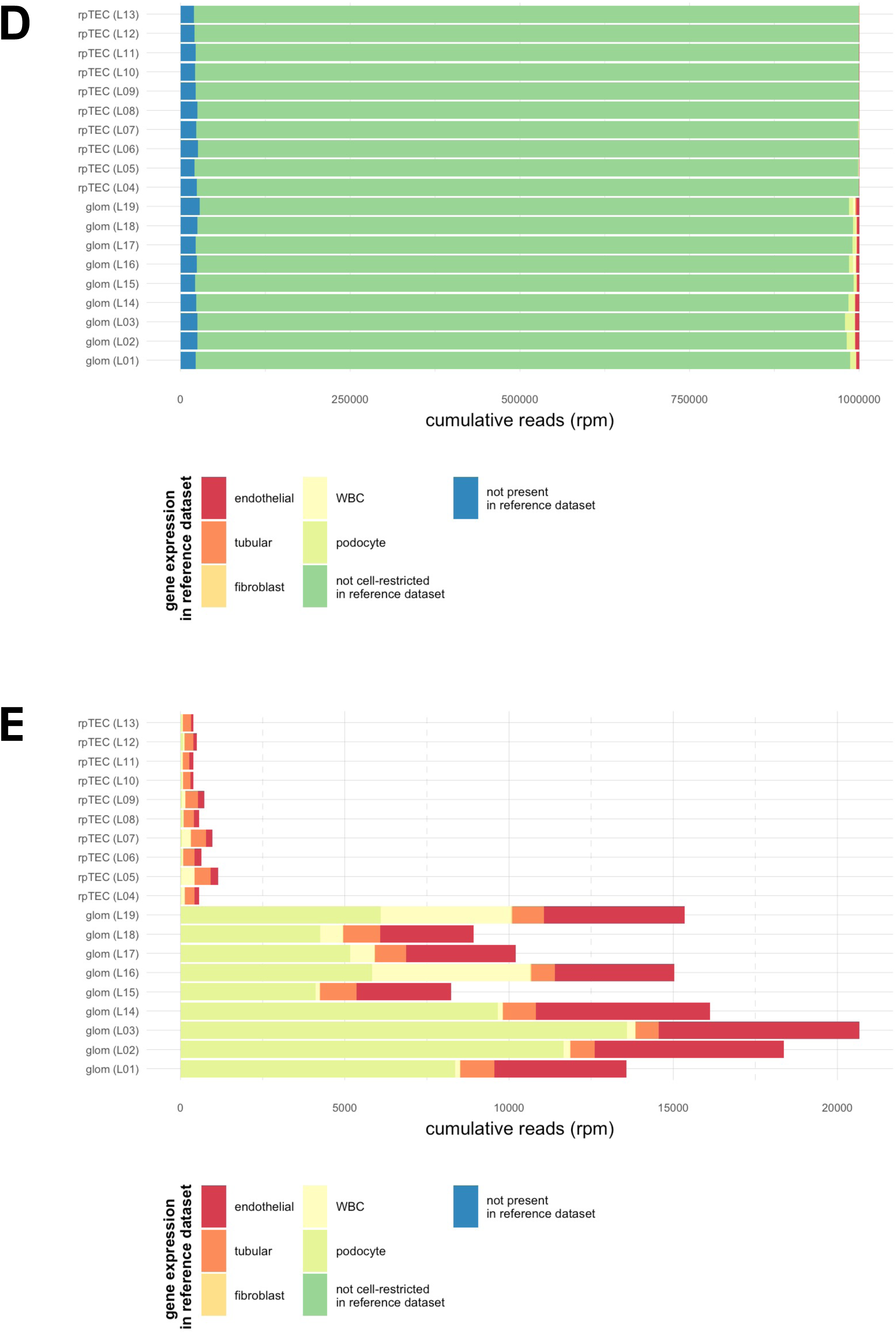
Validation of RNA alkylation and glomerular / tubular sample preparation. **A) Validation of glomerular preparations by immunoblot.** Whole kidneys or glomeruli were used to prepare protein samples for immunoblotting. Kidneys were obtained from podocin-UPRT mice (denoted Cre+) or Cre-negative littermate controls. The lane marked “L” was used to run the ladder; pencil marks at some ladder bands are visible. 20 micrograms of protein loaded per lane. After blotting the membrane was cut horizontally at ∼90 kDa; the upper portion was probed with anti-nephrin (expected band at 185 kDa) and the lower portion probed with anti-βactin (expected band at 45 kDa). **B) Alkylation of 4TU by iodoacetamide.** 1 mM 4-thiouracil controls were included alongside RNA samples in the alkylation reaction. UV spectrophotometry demonstrated the expected shift in absorbance, confirming successful alkylation of 4TU. **C) Glomerular and rTEC transcriptomes: unsupervised analysis.** In order to validate our protocols for isolating glomeruli and rTECs, we analysed the transcriptomes sequenced from glomerular and rTEC samples. In unsupervised hierarchical clustering and principal component analyses, glomerular and tubular libraries formed distinct clusters. **D)** & **E) Glomerular and rTEC transcriptomes: expression of known “marker” genes.** We used publicly-accessibly scRNAseq data to define “marker genes”, enriched in different kidney cell types. **D)** The majority of genes in our data showed uniform expression in this reference dataset (i.e. they were not cell-restricted). **E)** Same data are plotted, excluding reads mapping to genes with uniform expression (or no expression) in the reference dataset.

## References

1. Merchant, M. L., Rood, I. M., Deegens, J. K. J. & Klein, J. B. Isolation and characterization of urinary extracellular vesicles: implications for biomarker discovery. Nat. Rev. Nephrol. 13, 731–749 (2017).

2. Hunter, R. W. Extracellular RNA in kidney disease: moving slowly but surely from bench to bedside. Clin. Sci. Lond. Engl. 1979 (2020) doi:10.1042/cs20201092.

3. Hunter, R. W., Dear, J. W. & Bailey, M. A. Chapter 11 - Exosomes in nephrology. in Exosomes (eds. Edelstein, L., Smythies, J., Quesenberry, P. & Noble, D.) 257–283 (Academic Press, 2020). doi:10.1016/B978-0-12-816053-4.00011-0.

4. Lindoso, R. S. et al. Extracellular vesicles released from mesenchymal stromal cells modulate miRNA in renal tubular cells and inhibit ATP depletion injury. Stem Cells Dev. 23, 1809–1819 (2014).

5. Wang, B. et al. Mesenchymal Stem Cells Deliver Exogenous MicroRNA-let7c via Exosomes to Attenuate Renal Fibrosis. Mol. Ther. J. Am. Soc. Gene Ther. 24, 1290–1301 (2016).

6. Cantaluppi, V. et al. Microvesicles derived from endothelial progenitor cells protect the kidney from ischemia-reperfusion injury by microRNA-dependent reprogramming of resident renal cells. Kidney Int. 82, 412–427 (2012).

7. Thomou, T. et al. Adipose-derived circulating miRNAs regulate gene expression in other tissues. Nature 542, 450–455 (2017).

8. Ying, W. et al. Adipose Tissue Macrophage-Derived Exosomal miRNAs Can Modulate In Vivo and In Vitro Insulin Sensitivity. Cell 171, 372–384.e12 (2017).

9. Ying, W. et al. MiR-690, an exosomal-derived miRNA from M2-polarized macrophages, improves insulin sensitivity in obese mice. Cell Metab. 33, 781–790.e5 (2021).

10. Fong, M. Y. et al. Breast-cancer-secreted miR-122 reprograms glucose metabolism in premetastatic niche to promote metastasis. Nat. Cell Biol. 17, 183–194 (2015).

11. Castaño, C., Mirasierra, M., Vallejo, M., Novials, A. & Párrizas, M. Delivery of muscle-derived exosomal miRNAs induced by HIIT improves insulin sensitivity through down-regulation of hepatic FoxO1 in mice. Proc. Natl. Acad. Sci. U. S. A. 117, 30335–30343 (2020).

12. Sharma, U. et al. Small RNAs Are Trafficked from the Epididymis to Developing Mammalian Sperm. Dev. Cell 46, 481–494.e6 (2018).

13. Zhang, Y. et al. Hypothalamic stem cells control ageing speed partly through exosomal miRNAs. Nature 548, 52–57 (2017).

14. Villarroya-Beltri, C. et al. Sumoylated hnRNPA2B1 controls the sorting of miRNAs into exosomes through binding to specific motifs. Nat. Commun. 4, 2980 (2013).

15. Hoshino, A. et al. Tumour exosome integrins determine organotropic metastasis. Nature 527, 329–335 (2015).

16. Bruno, S. et al. Mesenchymal stem cell-derived microvesicles protect against acute tubular injury. J. Am. Soc. Nephrol. JASN 20, 1053–1067 (2009).

17. de Jong, O. G. et al. A CRISPR-Cas9-based reporter system for single-cell detection of extracellular vesicle-mediated functional transfer of RNA. 13.

18. Street, J. M. et al. Exosomal transmission of functional aquaporin 2 in kidney cortical collecting duct cells. J. Physiol. 589, 6119–6127 (2011).

19. Oosthuyzen, W. et al. Vasopressin Regulates Extracellular Vesicle Uptake by Kidney Collecting Duct Cells. J. Am. Soc. Nephrol. JASN (2016) doi:10.1681/ASN.2015050568.

20. Chevillet, J. R. et al. Quantitative and stoichiometric analysis of the microRNA content of exosomes. Proc. Natl. Acad. Sci. U. S. A. 111, 14888–14893 (2014).

21. Heil, F. et al. Species-specific recognition of single-stranded RNA via toll-like receptor 7 and 8. Science 303, 1526–1529 (2004).

22. Vallon, V. The proximal tubule in the pathophysiology of the diabetic kidney. Am. J. Physiol.-Regul. Integr. Comp. Physiol. 300, R1009–R1022 (2011).

23. Tang, S. C. W., Leung, J. C. K., Chan, L. Y. Y., Tsang, A. W. L. & Lai, K. N. Activation of Tubular Epithelial Cells in Diabetic Nephropathy and the Role of the Peroxisome Proliferator–Activated Receptor-γ Agonist. J. Am. Soc. Nephrol. 17, 1633–1643 (2006).

24. Kogot-Levin, A. et al. Proximal Tubule mTORC1 Is a Central Player in the Pathophysiology of Diabetic Nephropathy and Its Correction by SGLT2 Inhibitors. Cell Rep. 32, 107954 (2020).

25. Tabatabaeifar, M. et al. An inducible mouse model of podocin-mutation-related nephrotic syndrome. PLOS ONE 12, e0186574 (2017).

26. Nishizono, R. et al. FSGS as an Adaptive Response to Growth-Induced Podocyte Stress. J. Am. Soc. Nephrol. 28, 2931–2945 (2017).

27. Zoja, C., Abbate, M. & Remuzzi, G. Progression of renal injury toward interstitial inflammation and glomerular sclerosis is dependent on abnormal protein filtration. Nephrol. Dial. Transplant. 30, 706–712 (2015).

28. Wickman, L. et al. Urine Podocyte mRNAs, Proteinuria, and Progression in Human Glomerular Diseases. J. Am. Soc. Nephrol. 24, 2081–2095 (2013).

29. Matsushima, W. et al. Sequencing cell-type-specific transcriptomes with SLAM-ITseq. Nat. Protoc. (2019) doi:10.1038/s41596-019-0179-x.

30. Gay, L. et al. Mouse TU tagging: a chemical/genetic intersectional method for purifying cell type-specific nascent RNA. Genes Dev. 27, 98–115 (2013).

31. Muzumdar, M. D., Tasic, B., Miyamichi, K., Li, L. & Luo, L. A global double-fluorescent Cre reporter mouse. genesis 45, 593–605 (2007).

32. Belteki, G. Conditional and inducible transgene expression in mice through the combinatorial use of Cre-mediated recombination and tetracycline induction. Nucleic Acids Res. 33, e51–e51 (2005).

33. Truett, G. E. et al. Preparation of PCR-Quality Mouse Genomic DNA with Hot Sodium Hydroxide and Tris (HotSHOT). BioTechniques 29, 52–54 (2000).

34. Takemoto, M. et al. A New Method for Large Scale Isolation of Kidney Glomeruli from Mice. Am. J. Pathol. 161, 799–805 (2002).

35. Murata, F. et al. Distribution of Glycoconjugates in the Kidney Studied by Use of Labeled Lectins. J. Histochem. Cytochem. 31, 139–144 (1983).

36. Kusaba, T., Lalli, M., Kramann, R., Kobayashi, A. & Humphreys, B. D. Differentiated kidney epithelial cells repair injured proximal tubule. Proc. Natl. Acad. Sci. U. S. A. 111, 1527–1532 (2014).

37. Herzog, V. A. et al. Thiol-linked alkylation of RNA to assess expression dynamics. Nat. Methods 14, 1198–1204 (2017).

38. Moll, P., Ante, M., Seitz, A. & Reda, T. QuantSeq 3′ mRNA sequencing for RNA quantification. Nat. Methods 11, i–iii (2014).

39. Neumann, T. et al. Quantification of experimentally induced nucleotide conversions in high-throughput sequencing datasets. BMC Bioinformatics 20, 258 (2019).

40. Park, J. et al. Single-cell transcriptomics of the mouse kidney reveals potential cellular targets of kidney disease. Science 360, 758–763 (2018).

41. Börner, U., Szász, G., Bablok, W. & Busch, E. W. [A specific fully enzymatic method for creatinine: reference values in serum (author’s transl)]. J. Clin. Chem. Clin. Biochem. Z. Klin. Chem. Klin. Biochem. 17, 679–682 (1979).

42. R Core Team. R: A Language and Environment for Statistical Computing. (R Foundation for Statistical Computing, 2020).

43. Wickham, H. et al. Welcome to the tidyverse. J. Open Source Softw. 4, 1686 (2019).

44. Wilke, C. ggridges: Ridgeline Plots in ‘ggplot2’. (2021).

45. Zeileis, A., Kleiber, C. & Jackman, S. Regression Models for Count Data in R. J. Stat. Softw. 27, 1–25 (2008).

46. Canty, A. & Ripley, B. boot: Bootstrap R (S-Plus) Functions. (2021).

47. Hahne, F. et al. flowCore: a Bioconductor package for high throughput flow cytometry. BMC Bioinformatics 10, 106 (2009).

48. RGLab/flowWorkspace. GitHub https://github.com/RGLab/flowWorkspace.

49. Liang, Z., Breman, A. M., Grimes, B. R. & Rosen, E. D. Identifying and genotyping transgene integration loci. Transgenic Res. 17, 979–983 (2008).

50. Stec, D. E., Morimoto, S. & Sigmund, C. D. Vectors for High-Level Expression of cDNAs Controlled by Tissue-Specific Promoters in Transgenic Mice. BioTechniques 31, 256–260 (2001).

51. Lee, V. W. & Harris, D. C. Adriamycin nephropathy: A model of focal segmental glomerulosclerosis: Adriamycin nephropathy. Nephrology 16, 30–38 (2011).

52. Grgic, I. et al. Discovery of new glomerular disease–relevant genes by translational profiling of podocytes in vivo. Kidney Int. 86, 1116–1129 (2014).

53. Murillo, O. D. et al. exRNA Atlas Analysis Reveals Distinct Extracellular RNA Cargo Types and Their Carriers Present across Human Biofluids. Cell 177, 463–477.e15 (2019).

54. Grange, C. et al. Biodistribution of mesenchymal stem cell-derived extracellular vesicles in a model of acute kidney injury monitored by optical imaging. Int. J. Mol. Med. 33, 1055–1063 (2014).

