## Supplemental Table S1 for "Extracellular RNA moves from the glomerulus to the renal tubule"

Transnetyx®
Genetic Monitoring

### Genetic Monitoring Test Results

Type: Monitoring
Strain Panel: C57BL/6 x 129 $_{v2}$
Customer: Robert Hunter, University of Edinburgh
Order: 26498
Date Generated: Monday, October 5, 2020
Strain of Interest: C57BL/6

| **Order 26498 Results** | | |
|---|---|---|
| Sample: | 1098714 | 1104858 |
| Markers per sample: | 120 | 120 |
| # of alleles analyzed: | 238 | 238 |
| Homozygous C57BL/6: | 99 | 99 |
| Heterozygous | 18 | 16 |
| # of C57BL/6 alleles: | 216 | 214 |
| C57BL/6 Percentage: | 90.8% | 89.9% |

| Chr. | Location | dbSNP | 1098714 | 1104858 |
|---|---|---|---|---|
| 1 | 6655964 | rs13475709 | B6 | B6 |
| 1 | 21942441 | rs3711079 | HET | B6 |
| 1 | 46830176 | rs3712347 | HET | B6 |
| 1 | 67355864 | rs3662752 | HET | HET |
| 1 | 85781598 | rs13475962 | B6 | B6 |
| 1 | 125274326 | rs3022832 | UND | B6 |
| 1 | 146452387 | rs3666525 | B6 | B6 |
| 1 | 168216635 | rs3714825 | B6 | B6 |
| 1 | 189850841 | rs13476298 | B6 | B6 |
| 2 | 11043342 | rs3698941 | B6 | B6 |
| 2 | 37657113 | rs3709811 | B6 | B6 |
| 2 | 55195721 | rs13476509 | B6 | B6 |
| 2 | 76509857 | rs13476586 | B6 | B6 |
| 2 | 103247353 | rs3656441 | B6 | B6 |
| 2 | 132597468 | rs13476778 | B6 | B6 |
| 2 | 159982657 | rs3022939 | B6 | B6 |
| 3 | 11297853 | rs3680834 | B6 | HET |
| 3 | 26233740 | rs13477025 | B6 | HET |
| 3 | 51676615 | rs3709395 | B6 | B6 |
| 3 | 87370527 | rs4224040 | B6 | HET |
| 3 | 109769327 | rs3712218 | B6 | B6 |
| 3 | 130729788 | rs3089257 | B6 | HET |
| 3 | 153670143 | rs3691107 | B6 | B6 |
| 4 | 6893505 | rs3674478 | B6 | B6 |
| 4 | 27441980 | rs3661592 | B6 | B6 |
| 4 | 47147482 | rs3654185 | B6 | B6 |
| 4 | 70913565 | rs3654162 | B6 | B6 |
| 4 | 111526651 | rs13477912 | B6 | B6 |
| 4 | 135371936 | rs3679734 | B6 | B6 |
| 4 | 155717880 | rs3680364 | B6 | B6 |
| 5 | 10671078 | rs3676096 | B6 | B6 |
| 5 | 25416413 | rs3664933 | B6 | B6 |
| 5 | 45053753 | rs3663092 | HET | HET |
| 5 | 66662082 | rs4225249 | HET | Other |
| 5 | 92291809 | rs3704889 | HET | HET |
| 5 | 137471207 | rs3141573 | B6 | B6 |
| 5 | 149795466 | rs3722801 | B6 | HET |
| 6 | 3466870 | rs3661828 | B6 | B6 |
| 6 | 17499924 | rs13478647 | B6 | B6 |

| | | | | |
|---|---|---|---|---|
| 6 | 54549374 | rs3655236 | B6 | B6 |
| 6 | 75395550 | rs3671401 | B6 | B6 |
| 6 | 94352524 | rs3667765 | B6 | B6 |
| 6 | 128010127 | rs3726801 | B6 | B6 |
| 6 | 147673082 | rs3023840 | B6 | B6 |
| 7 | 4845207 | rs4226386 | B6 | B6 |
| 7 | 30723088 | rs3662246 | B6 | B6 |
| 7 | 44496336 | rs3710949 | B6 | HET |
| 7 | 75705579 | rs3679035 | C57BL/6\|129P\|129X1 | C57BL/6\|129P\|129X1 |
| 7 | 103468334 | rs3706526 | B6 | B6 |
| 7 | 139788374 | rs3700241 | C57BL/6\|129X1 | C57BL/6\|129X1 |
| 7 | 144896765 | rs4226997 | B6 | B6 |
| 8 | 15191287 | rs3709624 | HET | 129\|C57BL/6N |
| 8 | 36069255 | rs3660534 | Other | Other |
| 8 | 51011320 | rs3726383 | HET | HET |
| 8 | 68083563 | rs3089230 | Other | Other |
| 8 | 85260543 | rs3703660 | HET | HET |
| 8 | 98278863 | rs3710112 | B6 | B6 |
| 8 | 122947693 | rs3693295 | HET | HET |
| 9 | 17784920 | rs3663821 | B6 | B6 |
| 9 | 33798345 | rs3023205 | B6 | B6 |
| 9 | 47788124 | rs3686686 | B6 | B6 |
| 9 | 65238490 | rs3718417 | HET | B6 |
| 9 | 81859840 | rs3673457 | HET | B6 |
| 9 | 123642539 | rs3706619 | B6 | B6 |
| 10 | 6490661 | rs3663844 | B6 | B6 |
| 10 | 19373181 | rs3696310 | B6 | B6 |
| 10 | 53743149 | rs3696307 | B6 | HET |
| 10 | 83014931 | rs13480672 | B6 | B6 |
| 10 | 103008495 | rs3716716 | HET | B6 |
| 10 | 128453132 | rs3719409 | B6 | B6 |
| 11 | 4508730 | rs3659787 | B6J | B6J |
| 11 | 14367378 | rs3685856 | B6 | B6 |
| 11 | 24486229 | rs3673413 | B6 | B6 |
| 11 | 58970014 | rs3023311 | B6 | B6 |
| 11 | 82863758 | rs3663879 | B6 | B6 |
| 11 | 116315626 | rs3675087 | B6 | UND |
| 12 | 9543137 | rs3689696 | B6 | B6 |
| 12 | 32097332 | rs13481380 | B6 | B6 |
| 12 | 63972884 | rs3657568 | B6 | B6 |
| 12 | 75996767 | rs3663221 | B6 | B6 |
| 12 | 99200894 | rs3673029 | B6 | B6 |

| | | | | |
|---|---|---|---|---|
| 12 | 116179294 | rs3023711 | B6 | HET |
| 13 | 16184499 | rs3701757 | B6 | HET |
| 13 | 38430801 | rs3659063 | B6 | B6 |
| 13 | 68879859 | rs3714056 | B6 | B6 |
| 13 | 87674302 | rs3666540 | B6 | B6 |
| 13 | 115447807 | rs3724755 | B6 | B6 |
| 13 | 119390734 | rs6397687 | B6 | B6 |
| 14 | 9760330 | rs3689508 | B6 | B6 |
| 14 | 25343320 | rs3682880 | B6 | B6 |
| 14 | 55021861 | rs3697794 | B6 | B6 |
| 14 | 78038402 | rs3693589 | B6 | B6 |
| 14 | 120944588 | rs4230603 | B6 | B6 |
| 14 | 123146197 | rs3685710 | B6 | B6 |
| 15 | 10134594 | rs13482427 | B6 | B6 |
| 15 | 31916279 | rs3023416 | B6 | B6 |
| 15 | 42825752 | rs3667271 | B6 | B6 |
| 15 | 57160486 | rs3702158 | B6J | B6J |
| 15 | 97711920 | rs3023427 | HET | C57BL/6\|129X1 |
| 15 | 100692698 | rs6221928 | HET | B6 |
| 16 | 5596392 | rs4153115 | B6 | B6 |
| 16 | 19883079 | rs4165081 | B6 | B6 |
| 16 | 45037051 | rs4180179 | B6 | B6 |
| 16 | 63792473 | rs4195412 | B6 | B6 |
| 16 | 95108047 | rs4221067 | B6 | B6 |
| 17 | 23779934 | rs4231353 | B6 | B6 |
| 17 | 40843403 | rs3690398 | B6 | B6 |
| 17 | 58195862 | rs3710084 | B6 | B6 |
| 17 | 69049502 | rs3657117 | B6 | B6 |
| 17 | 86809846 | rs4231722 | B6 | HET |
| 18 | 5088196 | rs3089544 | HET | B6 |
| 18 | 21940443 | rs3707236 | HET | B6 |
| 18 | 54561772 | rs3715080 | HET | B6 |
| 18 | 77734915 | rs3725940 | B6 | B6 |
| 18 | 89459800 | rs3663208 | B6 | B6 |
| 19 | 18594885 | rs3691881 | C57BL/6\|129X1 | C57BL/6\|129X1 |
| 19 | 29265226 | rs3679607 | C57BL/6\|129X1 | C57BL/6\|129X1 |
| 19 | 38442814 | rs3695591 | C57BL/6\|129X1 | C57BL/6\|129X1 |
| 19 | 48137866 | rs3023496 | C57BL/6\|129X1 | C57BL/6\|129X1 |
| 19 | 60757991 | rs13483700 | B6 | B6 |
